# Revisiting the genetics of Lake Constance Coregonids using lake-wide whole genome sequencing

**DOI:** 10.64898/2026.01.18.700192

**Authors:** Arne Jacobs, Samuel Roch, Barnaby Roberts, Maria Capstick, Alexander Brinker

## Abstract

Anthropogenic pressures can have detrimental impacts on fish populations, with their effective management and conservation requiring accurate monitoring tools. Yet, this is not straightforward for closely-related, co-existing species that are difficult to distinguish using simple phenotypic or genetic approaches. Coregonids are of cultural and economic importance across Europe but have faced a multitude of pressures over the last century. Yet genomic management tools are lacking. In Lake Constance, a large pre-alpine lake, stocks have drastically collapsed due to a multitude of pressures, leading to a fishery closure. Here, we adopt a cost-effective, whole genome sequencing approach for lake-wide assessment of stock composition, spatial distribution and genetic diversity of highly admixed Lake Constance whitefish (*Coregonus* spp.). By sequencing 983 adult and larval genomes, we show that nearly 90% of the stock is made up by one of three species, the Gangfisch (*C. macrophthalmus*), and define the genetic relationship between Upper and Lower Lake Constance whitefish stocks. We also identified strong mixing between Gangfisch and Blaufelchen (*C. wartmanni*) on traditionally specific-specific spawning grounds, and detected strong admixture in larvae, with potentially drastic impacts on the effectiveness of hatchery supplementation and stocking. Despite the collapse and admixture, species still exhibit low to moderate levels of genetic diversity, maintain ecologically-relevant genetic differences, and seem to show differences in habitat use. Overall, we present a cost-effective, translatable tool for stock-wide sequencing and genetically-informed fisheries management, with our results calling for the re-evaluation of current management practices to avoid the potential genetic mixing between species.

## Background

Anthropogenic pressures, such as invasive species, eutrophication, and overfishing have strongly impacted fish populations worldwide, leading to broad ecological changes (Kefford et al., 2023; Reid et al., 2019; Tickner et al., 2020; Vardakas et al., 2025). These include population size collapses (Haase et al., 2025; Han et al., 2025), genetic diversity loss (Pinsky & Palumbi, 2014), increased hybridization (Vonlanthen et al., 2012), and altered habitat use (Alexander et al., 2017). Understanding the impacts of anthropogenic pressures on fish populations requires large-scale monitoring, yet this can be highly challenging when multiple species or ecotypes co-exist that cannot be reliably phenotypically distinguished.

Fisheries management often relies on targeted genotyping assays (e.g. GT-seq, microsatellites) and DNA barcoding for genetic monitoring (Bernatchez et al., 2017; Friedman et al., 2022; Theissinger et al., 2023). While these approaches can be powerful and cost-effective once developed, their development is often expensive and labour-intensive, they only cover a small portion of the genome (few microsatellites to few hundred SNPs) and can be prone to ascertainment bias if they have not been developed across all relevant populations (Beemelmanns et al., 2024; Robledo et al., 2018). Whole-genome sequencing (WGS) offers a powerful approach for investigating fine-scale population structure and differentiation for conservation and fisheries management (Bernatchez et al., 2017; Leitwein et al., 2020; Tengstedt et al., 2025). Yet, WGS is often prohibitively expensive and requires large quantities of DNA, limiting its suitability in many applied contexts. The advent of low-coverage whole genome sequencing (‘lcWGS’) opens up exciting opportunities to rapidly and cost-effectively sequence the entire genome of an individual from relatively little DNA (Gaio et al., 2022; Lou et al., 2021; Therkildsen & Palumbi, 2017). In contrast to traditional approaches, lcWGS can be used to simultaneously investigate population structure, perform stock/species assignment and also investigate fine-scale patterns of genetic differentiation and selection across the genome for a large number of individuals, even when levels of genetic differentiation is weak (DeSaix et al., 2024; Lou et al., 2021; Therkildsen & Palumbi, 2017). However, it has been rarely applied at a large scale to directly inform fisheries management decisions.

European whitefish (*Coregonus spp*.) are economically and culturally important fisheries species in Europe, including in Lake Constance, one of the largest pre-alpine lakes in Central Europe (Haase et al., 2025; Vonlanthen et al., 2012). Originally, Upper Lake Constance (ULC) harboured four ecologically distinct whitefish species; the pelagic spawning, planktivorous Blaufelchen (*C. warmanni*), the generalist pelagic-littoral Gangfisch that spawned in the benthic zone (*C. macrophthalmus*), the shallow-spawning littoral Sandfelchen (*C. arenicolus*), and the now-extinct profundal Kilch (*C. gutturosus*) (Steinmann, 1950; Vonlanthen et al., 2012). Furthermore, the Lower Lake Constance (LLC), a smaller and shallower lake-basin that is connected to Lake Constance through a short river, contained the Unterseefelchen (or Weissfelchen), which has not been genetically characterised (De-Kayne et al., 2022; Steinmann, 1950). Anthropogenic eutrophication in the 20th century led to extinction of the profundal Kilch and the collapse of reproductive barriers between species, thereby increasing hybridization by shifting spawning and feeding habits (Alexander et al., 2017; Feulner & Seehausen, 2019; Frei, Mwaiko, et al., 2023; Hirsch et al., 2013; Jacobs et al., 2019; Numann, 1978, 1986; Schweizer, 1894; Vonlanthen et al., 2012). This led to reduced genetic differentiation between species in Lake Constance, particularly between the pelagic Blaufelchen and the Gangfisch (Frei, Mwaiko, et al., 2023). Although hybridization led to increased allelic richness within species (Gum et al., 2014; Vonlanthen et al., 2012), an overall loss in genetic diversity was observed in all whitefish species over the last century (Frei, Mwaiko, et al., 2023). Over the last decades, a combination of overfishing, environmental change, and invasive species (Baer, Spiessl, et al., 2022; Dahms et al., 2024; Haase et al., 2025; Rösch et al., 2018) have contributed to a collapse of the whitefish populations, especially the Blaufelchen, a key fisheries target (Haase et al., 2025), leading to the closure of the commercial whitefish fishery in Lake Constance in 2024 (Baer et al., 2016; Haase et al., 2025). Despite extensive stocking over the last 150 years, larval recruitment has been low (Baer et al., 2023; Haase et al., 2025), raising questions on the efficiency of stocking efforts. To better understand the current situation and inform effective management strategies, more detailed information on the contemporary composition and habitat use of European whitefish species across all of Lake Constance, from larvae to adults, is required.

Fisheries management strategies often rely on historical knowledge of species, including their genetic relationships, levels of genetic diversity and habitat preferences. Yet, these factors can rapidly change in the face of environmental and population change. Here, we integrate large-scale sampling and phenotypic analysis of European whitefish in Upper and Lower Lake Constance across several years, seasons and developmental stages with novel, cost-effective low-coverage whole genome sequencing (lcWGS) of nearly 1,000 individuals to reconstruct the contemporary genetic composition of European whitefish in Lake Constance, investigate the spatial distribution of individual species, and map genomic levels of genetic diversity and differentiation across species. Using this comprehensive approach, we demonstrate the suitability of lcWGS for large-scale cost-effective fisheries management and show that historical patterns of species distribution are not reliable indicators of species composition, identify the mixing of species on spawning grounds, genetically identify Unterseefelchen as Sandfelchen (*C. arenicolus*), and highlight low levels of genetic diversity and differentiation. Thus, these results question the direction and effectiveness of current management strategies.

## Methods

### Sampling

Adult individuals of European whitefish species occurring in Lake Constance were targeted using benthic and pelagic gillnets during sampling campaigns from 2021 to 2023 (Table S1, Fig. 1). We targeted all three species in the Upper Lake Constance (ULC), sampling “Blaufelchen” (*C. wartmanii*) and “Gangfisch” (*C. macrophthalmus*) during the annual spawning fisheries in the pelagic and littoral zone, respectively. The littoral “Sandfelchen” (*C. arenicolus*) was repeatedly targeted and sampled over a longer timeperiod in the littoral zone of the lake. Whitefish from the Lower Lake Constance (“Unterseefelchen” in LLC) were sampled in various locations. Whitefish that migrate in September and October into the Alpine Rhine to spawn (“Alpenrheinfelchen”) were targeted by local anglers (n=1 sample). Furthermore, adult whitefish were caught as part of a lake-wide fishing campaign in September 2024, during which the entire lake was systematically fished using 391 randomized benthic and pelagic gillnets in varying water depths (Fig. 1; Vonlanthen et al., in preparation). A total of 34 whitefish in the Lower Lake and 70 whitefish in the Upper Lake were caught (Table 1 and S2). Fish were stunned with a blow on the head, killed with a cut at the gills and total length (0.1 cm) and wet weight (0.1 g) were measured. All individuals from the initial fishing campaign and most individuals from the lake-wide fishing campaign were photographed laterally using a digital camera (Pentax K3 II with 18–135 mm lens and fixed focal length). For genetic analysis, fin clips (∼0.5 cm^2^) were preserved in absolute ethanol.

**Figure 1.**
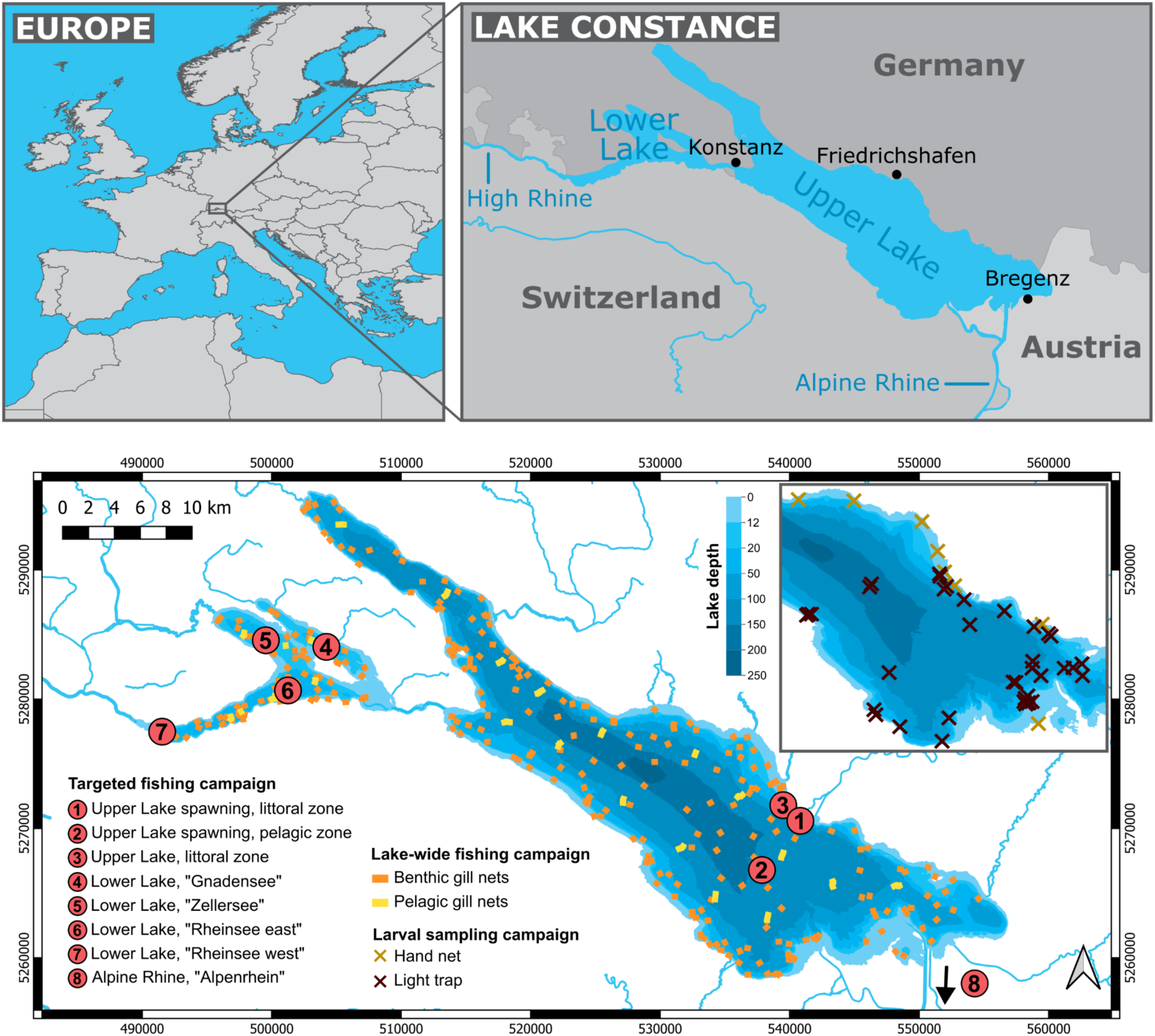
Sampling information. Overview over the study area and sampling locations for adult and larval whitefish in Lake Constance, southwest Germany. The lower relief map shows the sampling locations for each fishing campaign, with explanations in the legend. Red dots in the main map show the locations of targeted fishing campaigns, whereas the yellow and orange squares show the locations of all benthic and pelagic gill nets used for the lake-wide fishing campaign. Crosses in the insert map show the sampling locations of larvae in Upper Lake Constance. More Details can be found in Tables S1-S3.

Larval whitefish were captured in the pelagic and littoral zone of ULC (Fig. 1) using light traps developed at the Fisheries Research Station. Larvae are attracted toward a pipe via a light and subsequently suctioned into the pipe and retained through a pump. The capture of larvae in the light traps is almost always non-lethal. In 2021 and 2022, five light-traps were deployed at >20 m depth. In 2022, five additional traps were deployed in the deeper littoral zone (∼5 m) of the lake. Light traps were deployed from February to May in both years. In the shallower littoral zone larvae were captured by hand in 2021/22, either on foot or onboard a boat between March and June in 2021 and 2022. Larvae were attracted using headlamps or spotlights and captured using aquarium hand-nets. A total of 744 larvae were caught (Table 1 and S3) and preserved in absolute ethanol.

All individuals included in the current study were caught by licensed personnel with permission of the local fisheries administration (Regierungspräsidium Tübingen) and according to German animal protection legislation (§4) and the ordinance on the slaughter and killing of animals (Tierschutzschlachtverordnung §13).

### Phenotypic analysis

To identify phenotypic differences between whitefish species in Lake Constance, we performed landmark-based body shape analyses and analyses of linear measurements (see Supplementary Methods for details). We compared body shape among the genetically identified whitefish species using 17 homologous landmarks (Fig. S1A, Table S5) using the *geomorph* V4.0.10 (Baken et al., 2021). Generalized Procrustes Analysis was performed to remove size, position and orientation differences, and individuals with abnormal body shape (e.g. due to body arching) were identified using the *plotOutliers* function and excluded if necessary, leaving 158 individuals (Table 1). We tested for differences in shape between species using a permutation-based procrustes ANOVA with residual randomization with the following model: coords ∼ log(size) * species. We did not perform a size correction, as allometric effects were different across species. Instead, we post-hoc compared species using the RRPP package with the null model: coords ∼ log(size) to account for allometric effects (Collyer & Adams, 2018) and adjusted p-values were adjusted using the Bonferroni-Holm method. Shape differences were illustrated using a PCA in *geomorph* and wireframe models. A canonical variate analysis (CVA) was used to examine the accuracy of species classification based on body shape.

Furthermore, we investigated phenotypic differences using 15 linear measurements for the same 158 individuals (Fig. S1B, Table S6) (see Supplementary methods). Highly correlated linear measurements (spearman rank coefficients >0.9) were excluded, and we tested different size correction approaches. We used Recursive Feature Elimination (RFE) to identify linear measurements that distinguished the species. Measurements with the highest classification accuracy were used for PCA in *MorphoTools2* V1.0.2.1 (Šlenker et al., 2022) and we performed a K-nearest-neighbor classificatory discriminant analysis to estimate the accuracy of linear measurements for species classification in *MorphoTools2*.

### Whole genome sequencing

DNA was extracted from fin clips for larval (n=744) and adult (n=239) samples using a magnetic bead-based protocol (dx.doi.org/10.17504/protocols.io.b46bqzan), concentrations quantified using a Qubit Flex (1X BR dsDNA assay), and quality determined using agarose gels. Whole genome sequencing (WGS) libraries were prepared using a modified HackFlex protocol (Gaio et al., 2022) (see Supplementary File 1 for protocol) from 10ng of DNA per individual. Compared to the original protocol, we reduced the amount of Dimethylformamide (DMF) in the tagmentation buffer to 20% from 50% to minimise the amount of this highly toxic chemical without impacting library quality, and we performed a double-sided bead clean up on the final libraries. Final libraries were quantified using the Qubit Flex (HS dsDNA assay) and the fragment size distribution for a subset was checked on the Tapestation (D5000 assay), with an expected peak around 400-600bp. We pooled libraries at equal molarities. We sequenced up to 288 libraries per pool using 150PE reads on NovaSeq X Plus 10B lanes at Novogene UK to an average of 1.6 Gb per individual. Some individuals were sequenced on an additional NovaSeq X Plus 25B lane to increase coverage. Furthermore, we downloaded public whole genome data for adult individuals with confirmed species assignment from ENA for Gangfish, Sandfelchen, and Blaufelchen (Frei et al. 2023, 2022) (Table S4).

### Bioinformatic processing and variant identification

Raw sequencing reads were processed following best practices for lcWGS data (Lou et al., 2021; Lou & Therkildsen, 2022) (see Supplementary methods for detailed parameters) using *fastp* (Chen et al., 2018). Processed reads were mapped to the European whitefish (‘Balchen’) reference genome (AWGv2; (De-Kayne & Zoller, 2020)) using *bwa mem* (Li & Durbin, 2009), replicated samples merged and duplicated reads removed using *sambamba* v.0.8.2 (Tarasov et al., 2015), and overlapping reads clipped using the *bamUtils clipOverlap* (https://github.com/statgen/bamUtil). Bam files were indexed using *sambamba* before indel realignment using *GATK3.8*. We estimated the depth of coverage per bam file using *mosdepth* v.0.3.11 (Pedersen & Quinlan, 2018).

Due to the low coverage of the sequencing data, we performed genotype likelihood-based (GL) analyses with *ANGSD* v0.938 (Korneliussen et al., 2014). We created two different SNP datasets to address distinct questions. For the first list, we identified SNPs across all 250 adult samples (‘adult SNP dataset’) using the samtools genotype likelihood model, a minimum SNP value of 1e-6 and determined major/minor alleles based on allele frequencies. We removed sites below a total sequencing depth of 250 (number of individuals x mean sequencing depth) and above 1300 (2 x number of individuals x mean sequencing depth), and removed sites with missing data in more than 20% of individuals, minimum base quality below 30, a minimum individual depth below 1, multi-mapping reads, unmapped or duplicated reads and those without a matching pair. We also adjusted the mapping quality for excessive mismatches and only kept SNPs with a minor allele frequency of 5%. Lastly, we removed SNPs falling within potentially problematic genomic regions that can impact analyses accuracy. We removed SNPs with excess heterozygosity potentially due to collapsed paralogs, estimated using the *ngsParalog calcLR* (Dallaire et al. 2023), excluded all sites within likely collapsed regions in assembly (Frei et al. 2023), and removed sites within regions without unique mappability (Wang et al. 2024), as inferred using *genmap* (with 150-mers) (Pockrandt et al. 2020).

For the second list, we created an additional SNP dataset for a subset of adult reference samples (published data and individuals from targeted sampling) that were used for species assignments of larvae (‘larval SNP dataset’). We identified variant sites across adult reference samples (Table S2) using the same filtering strategy as above using adjusted filtering values. Since larvae had ultra-low sequencing coverage, we did not use them for SNP identification but instead inferred genotype likelihoods for larvae based on the filtered ‘larval SNP dataset’ with the -sites function We did not apply any other filters at this step, such as depth filters, due to the very low coverage (DeSaix et al. 2024).

### Population structure

We investigated population structure across adults using PCAngsd (Meisner & Albrechtsen, 2018), using published reference individuals to identify genetic clusters corresponding to species, refine the species-assignment of adults caught during the targeted fishing campaign based on their clustering PCA space, and assign individuals caught across the upper and lower lake in 2024 to their respective species. Across adult and larval samples, we estimated the pairwise IBS matrix across all individuals based on randomly sampled single reads for each site (-doIBS 1) for MDS analysis. The random sampling approach is less biased to variation in coverage across samples (Lou et al., 2021) and better suited for samples with ultra-low coverage and varying sequencing depth. Processing and plotting of PCA and MDS results were performed in R.

We investigated the correlation between morphology (based on landmarks and linear measurements) with genetic variation to determine if genetically intermediate individuals are also phenotypically intermediate. We used composite principal component scores across PC1 and PC2 as a measure for genetic and phenotypic variation (Verta & Jones, 2019), which we estimated for the genetic PCA, landmark-based PCA and linear-measurement PCA independently, by summing up PC1 and PC2, weighting each by their eigenvalues. We used linear mixed models (*lmer* in the lme4 R-package) to estimate the correlation between genetic variation and phenotypic variation, with species as a random factor.

### Larval Species assignment

We assigned larvae to their respective species using a probabilistic framework in *WGSassign* (DeSaix et al. 2024). Assignment accuracy is heavily biased by differences in sample size and sequencing depth between reference populations (DeSaix et al., 2023, 2024). To minimise biases, we normalised the effective sample size (ESS) of reference populations by subsampling an equal number of adult individuals per species with similar sequencing depths (DeSaix et al., 2024). We tested our assignment accuracy based on adult samples of known origin that were not part of the reference populations and used *sambamba view* to subsample four test samples to mean and minimum sequencing depths observed for larval samples (0.5X, 0.1X and 0.01X) to test the effect of sequencing depth on assignment accuracy. As we had 100% accuracy for downsampled samples, we assumed similar assignment accuracy for larvae. Since admixed individuals have mosaic genomes that cannot be completely assigned to one species, and because posterior probabilities of assignment are suggested to be unreliable for lcWGS data, we split up the SNP dataset into 10 equal sets of ∼240,000 SNPs, performed the species assignment for each subset separately, to estimate the consistency of assignment across sections of the genome (DeSaix et al., 2023, 2024). We determined an assignment as accurate if 8 of 10 genomic subsets (0.8) assigned an individual to the same species.

### Population genomics adults

To investigate the contemporary genomic diversity of whitefish in Lake Constance, we performed GL-based population genomic analyses in adult whitefish. First, we investigated patterns of Linkage disequilibrium (LD) decay across the genome by species using *ngsLD* (Fox et al., 2019). We estimated LD between all SNPs within 4Mb and used these LD estimates to infer patterns of LD decay and reconstruct recent population trends over the last few hundred generations in *SNeP* (Barbato et al., 2015). Second, we inferred genome-wide genetic diversity (nucleotide diversity [π] and Tajima’s D) across all filtered sites (variant and invariant) using *ANGSD*, compared levels of π between species using an ANOVA and Tukey’s Posthoc test in R, and tested if levels of Tajima’s D were different from zero using a Wilcoxon rank sum test. Third, we inferred the landscape of differentiation and selection between species across the genome. We estimated Fst between all species pairs for SNPs and in 50kb sliding windows with 25kb steps (based on LD decay rates) using *ANGSD* and *realSFS*. We defined outlier regions as Fst values above the 99% of the Fst distribution. Furthermore, we estimated the population branch excess statistic (PBE) for each species from windowed Fst data (see Supplementary methods for details). We identified outlier windows as those with PBE values in the top 0.1% of the genome-wide distribution. Lastly, we identified genomic outlier regions indicative of low recombination regions and structural variants using a modified *LOSTRUCT* approach (Li & Ralph, 2019) (see Supplementary Methods).

## Results

### Population structure and species composition

Using the adopted HackFlex protocol, we cost-effectively generated low whole-genome sequencing data for 239 adult whitefish (mean depth of coverage = 2.6X; range 0.01X to 8.83X; Fig.S2), moderate PCR-duplications rate (mean=8%, range=1.7%-11.1%), and identified 4,206,794 filtered SNPs across all adults. The PCA separated the three species in ULC into three clusters along PC1 and PC2 (Fig.2A, Fig.S3-S4). However, 15 adults that caught during the targeted campaign on spawning grounds were assigned to the wrong species in the field, with 13 Blaufelchen assigned as Gangfisch and two Gangfisch as Blaufelchen (Fig.S3). Species assignments were corrected based on the genomic data. Unterseefelchen from Lower Lake Constance were genetically indistinguishable from Sandfelchen. The single individual caught in the rhine inflow (‘Alpenhein’) genetically clustered with Sandfelchen in PCA space. Furthermore, the species assignment of the 105 adults caught during the lake-wide survey in 2024 identified 7 Blaufelchen (6.7%% of individuals) and 63 Gangfisch (60%). As we couldn’t distinguish Sandfelchen and Unterseefelchen, we assigned individuals from that genetic cluster caught in ULC as Sandfelchen (12; 11.4%) and those caught in LLC as Unterseefelchen (23; 21.9%).

**Figure 2.**
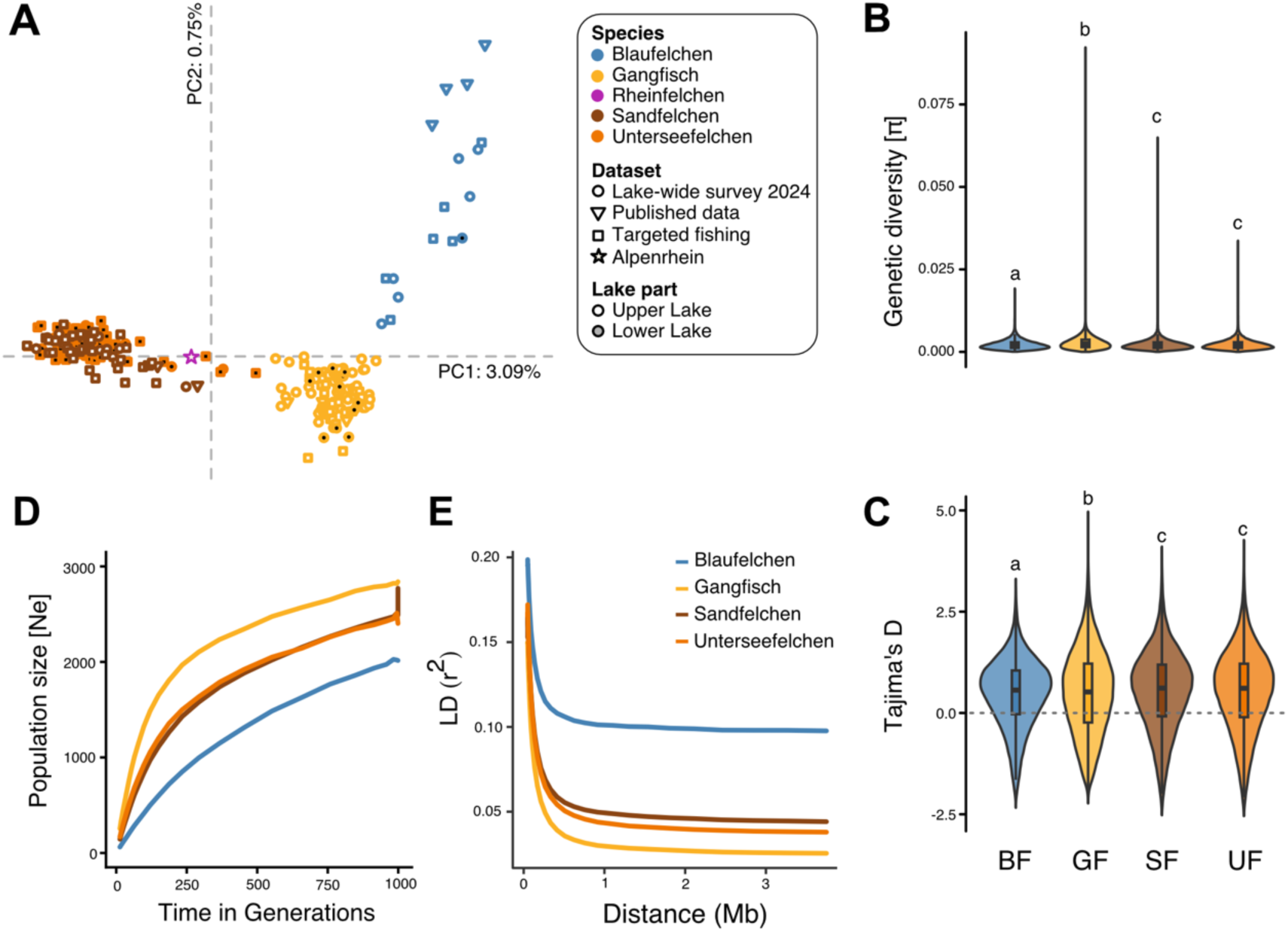
Genetic diversity and population history. (A) Principal components analysis showing the population structure of adult whitefish in Lake Constance. Individuals are coloured by species and shapes are filled in by catch location. The shape represents the dataset. See legend for details. (**B, C**) Nucleotide diversity (B) and Tajima’s D (C) differed between species in Upper Lake Constance but not between Sandfelchen and Unterseefelchen. Statistically different groups are highlighted by letters above the violin plots. See text for statistical results. (**D**) Changes in effective population size (Ne) over the last 1,000 generations by species as estimated from linkage disequilibrium (LD) data with SNeP. (**E**) LD decay by species as estimated with SNeP.

### Low genetic diversity and population declines

Estimates of genetic diversity and population history indicated low levels of genetic diversity across all species, with significant differences between species (ANOVA: F_(3)_=1697, p<2e-16), and values of π ranging from 0.00230 Blaufelchen to 0.00234 in Sandfelchen and 0.00237 Unterseefelchen and 0.00282 in Gangfisch (Fig.2B). Genome-wide levels of Tajima’s D were skewed toward positive values in all populations and statistically different from zero (Wilcox test, p<0.05), indicative of sudden population contraction (Fig. 2C).

LD-based analyses indicated declines in effective population sizes (‘Ne’) over the last 1,000 generations in all species (Fig. 2D,E). Blaufelchen showed the lowest contemporary Ne of 62, the lowest levels of genetic diversity and highest levels of LD (Fig. 2D,E). In contrast, Sandfelchen and Unterseefelchen had estimated Ne’s of 144 and 169, respectively, and Gangfisch had the highest estimated Ne of 255. While Blaufelchen showed a continuous steep decline in Ne over the last 1,000 generations, the other species showed steeper declines in Ne in the last 100 - 200 generations (Fig.2D).

### Landscape of genetic differentiation and selection

Genome-wide pairwise Fst between species was overall low, ranging from 0.006 between Sandfelchen and Unterseefelchen to 0.043 between Blaufelchen and Unterseefelchen (Fig. 3A, Fig. S5, Table S7). Genome-wide scans of differentiation revealed peaks of elevated Fst between species across the genome (Fig.2A, Fig.S5), although overall Fst values were still low (< 0.5). Comparing outlier Fst windows (Fst > 0.99%) to known adaptive trait QTL, we detected elevated differentiation Gangfisch and Sandfelchen around the *edar* gene on chr23, which has been associated with variation in gill raker count in the Alpine whitefish radiation (Fig.3B) (De-Kayne et al., 2022). These species are known to differ in gill raker count, with Sandfelchen having fewer gill rakers than the other species (Hirsch et al., 2013; Vonlanthen et al., 2012). The same region did not show strong differentiation between the other whitefish species but showed signatures of selection in Blaufelchen based on population branch excess (PBE > 0.999%; Fig.3B, Fig.S6). Genomic signatures of selection across the genome were species-specific, without shared outlier windows (PBE > 0.999%) across species (Fig.S6). Furthermore, local PCA identified 83 outlier regions across the genome (>4 SD from the mean per MDS axis), which showed distinct patterns of population structure compared to the rest of the genome, potentially due to reduced recombination rates and/or structural genomic variation (Li & Ralph, 2019) (Fig. S7 and S8). Of these local PCA outlier windows, between 31 and 143 overlapped with Fst outlier windows in at least one comparison, including the Fst outlier region on chr23 associated with gill raker count. Most outlier regions showed PCA patterns indicative of low-recombination regions, i.e. without three distinct clusters, except the outlier regions on chr8 and chr24 (Fig. S9) (Mérot et al., 2021). However, these regions did not distinguish species and did not overlap Fst outlier regions.

**Figure 3.**
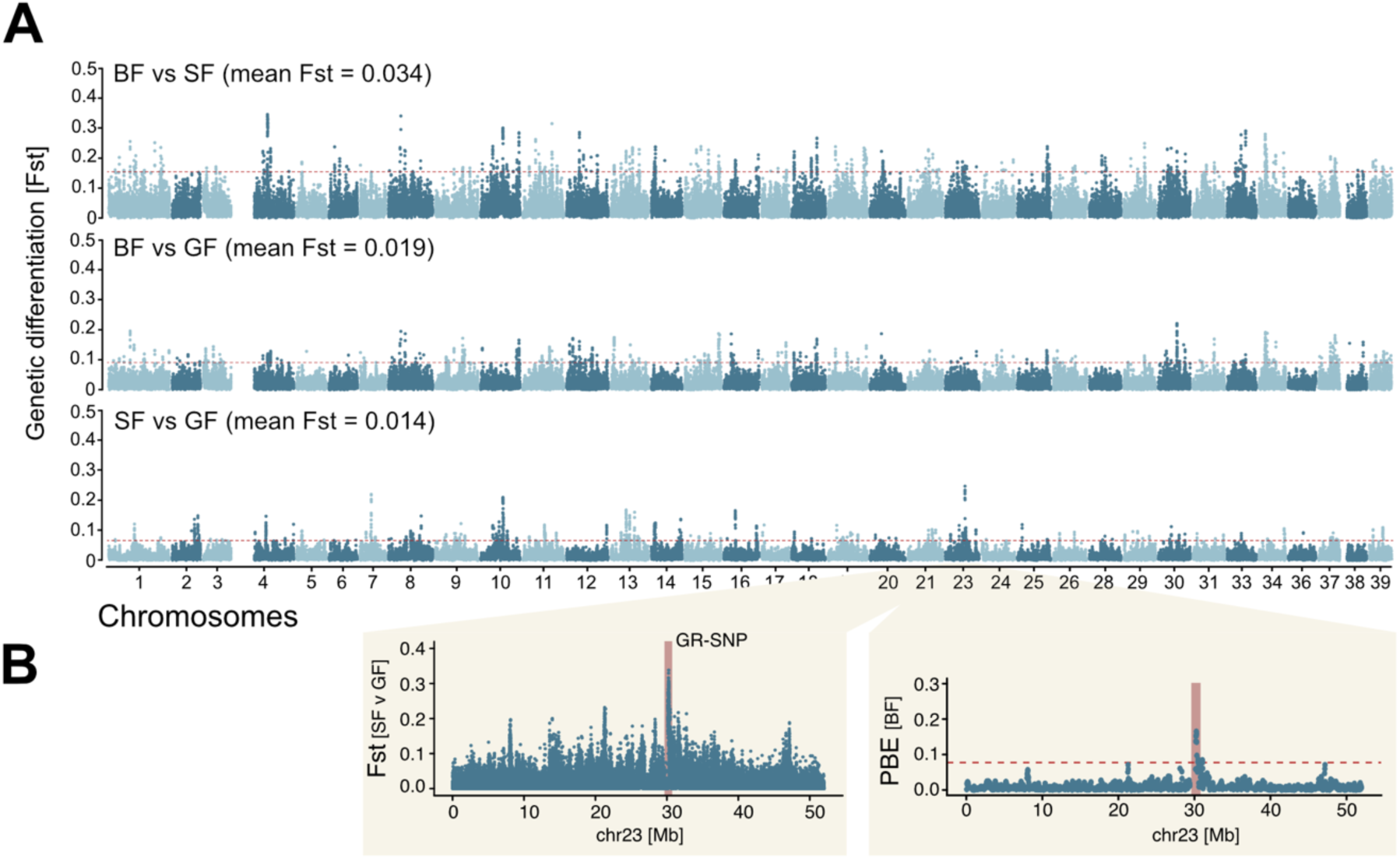
Genetic differentiation. (**A**) Manhattan plots showing the levels of genetic differentiation in 50kb sliding windows (25kb steps) across the genome between species in Upper Lake Constance. SF = Sandfelchen; BF = Blaufelchen; GF = Gangfisch. (**B**) The plot on the left shows SNP-based Fst along chr23, highlighting increased differentiation between SF and GF around a SNP previously associated with variation in gill raker count in European whitefish. The plot on the right shows the PBE value in Blaufelchen for the same genomic region, with the gill raker count associated region highlighted in red.

### Phenotypic results

We assessed phenotypic variation between adult whitefish based on genetic species assignment (Table S8). Body size had a statistically significant, species-dependent effect on body shape based on landmarks (Z [effect size] = 1.790, p = 0.038), with shape differing between species (Z = 1.687, p = 0.046). Shape differed between Blaufelchen and Sandfelchen (Z = 2.528, p = 0.024), Blaufelchen and Unterseefelchen (Z = 4.014, p = 0.001), and Gangfisch and Unterseefelchen (Z = 3.187, p = 0.006) (Fig.4A). However, there was considerable overlap in shape between species (Fig.4A), with changes in body shape along PC1 primarily being explained by the degree of ventral curvature and head position and changes along PC2 mainly being confined to body height and head length (Fig. 4B). Overall classification accuracy was 65.8% (Table S7), with the highest accuracy for Unterseefelchen (75.0%) and the lowest for Blaufelchen (33.3%).

**Figure 4.**
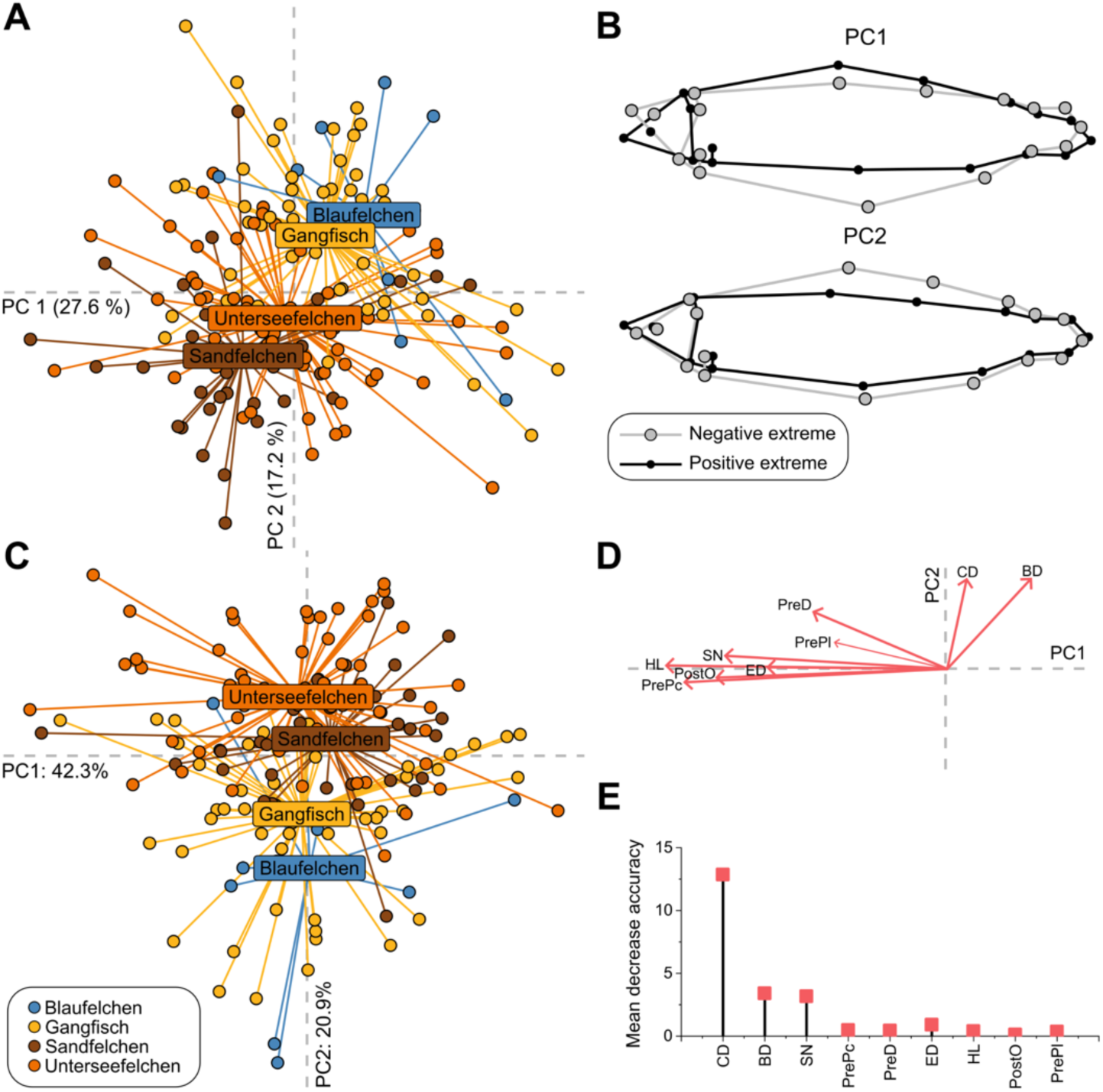
Phenotypic variation. (**A**) Scatter plot showing the first two principal components (PCs) of the phenotypic variation of identified whitefish species, based on 17 homologous landmarks. The first two PCs are explaining 44.8% of the variance of the data. Each point represents one individual, which is coloured based on their genetic assignment. (**B**) Wireframe graphs of the shape comparing the extremes along the PC axes in the landmark PCA. (**C**) Scatter plot showing the first two principal components (PCs) of the phenotypic variation of identified whitefish species, based on selected linear measurements. The first two PCs are explaining 63.2% of the variance of the data. (**D**) Loading plot for the linear measurement PCA. (**E**) Mean decrease accuracy values based on a Random Forest model, showing importance of selected linear measurements for species differentiation.

For linear measurements, we excluded PostA and PostD due to high correlations of PreA and PostA (r = -0.91) and PreD and PostD (r = -0.94) (Fig.4C,D) and we identified the following CD, BD, SN, PrePc, PreD, ED, HL, PostO, PrePl as optimal features for distinguishing whitefish species (Fig.4E). For the PCA, we only used the selected measurements listed above, and similar to the landmark data, species showed a considerable overlap in phenotype (Fig. 4C). Species separated largely along PC1, which was explained by variation in Caudal peduncle depth (CD) and body depth (BD) (Fig. 4D). The importance of CD and BD for species differentiation was supported by mean decrease accuracy values derived from the RFE (Fig. 4D). Overall classification accuracy based on K nearest neighbour classificatory discriminant analysis (K = 7) was 62.7% (Table S9), with the highest accuracy again for Unterseefelchen (83.6%) and lowest for Blaufelchen (0%).

Phenotypic variation was significantly correlated with genetic variation (compound PC1 + PC2) for landmarks (Linear mixed model: beta = 1.46, 95% CI [0.70, 2.22], t(137) = 3.81, p < .001), but not for linear measurements (Linear mixed model: beta = -22.46, 95% CI [-103.70, 58.78], t(137) = -0.55, p = 0.585) (Fig.S10). Thus, overall body shape seems to vary with genetic variation, with admixture potentially leading to intermediate body shapes.

### Species assignment of whitefish larvae

We sequenced 744 larvae to a mean depth of 0.5X (range 0.01X to 3.55X; Fig.5A) and inferred genotype likelihoods at a maximum of 2,446,925 SNPs Due to the low sequencing depth and breadth, larvae had on average sufficient sequencing data at 38% of all SNPs (ranging from 0.7% to 95%), which is sufficient for inferring population structure and population assignment (DeSaix et al., 2024). In comparison, adults covered ∼90% of all sites. Standardisation of effective sample size (ESS) for each reference group resulted in 10 individuals per species, with ESS from 6.5 to 6.9 (Fig.5B). The assignment accuracy of test samples of known species was 100% for all subsampled coverages (0.5x, 0.1x and 0.01x) with 100% assignment consistency across all ten SNP subsets, indicating sufficient power to accurately assign larvae to their species down to 0.01x coverage. Species assignment of larvae using lcWGS showed that 91.3% of individuals were Gangfisch (679 out of 744), with 7.3% being assigned as Blaufelchen (54 out of 744), and only 1.5% were assigned as Sandfelchen (11 out of 744) (Fig.5B). Gangfisch had a marginally lower ESS (6.5) compared to Blaufelchen (6.9) and Sandfelchen (6.8), suggesting that this large assignment bias between species is not driven by differences in ESS <u>(DeSaix et al. 2023)</u>. Three individuals were not consistently assigned to the same species, with assignment consistencies below 0.8. Of these, two individuals showed assignment consistencies of 0.5 to Gangfish and Blaufelchen, suggesting recent admixture of these individuals. The third individual showed an assignment consistency of 0.7 to Gangfisch and 0.3 to Blaufelchen. These three individuals were also intermediate in their position in the MDS plot between Blaufelchen and Gangfisch, suggesting that these are potentially recent hybrids (Fig.5A). Like the species assignment, most larvae clustered genetically with adult Gangfisch in the MDS plot (Fig.5A). Larval whitefish showed a broader spread in the genomic space compared to adults, with many individuals clustering intermediate between species, especially Gangfisch and Blaufelchen. While this could potentially be explained by the lower coverage for many samples, we found that downsampled test samples with coverages down to 0.01x still clustered closely with the other adult samples (Fig.S11), suggesting that coverage alone does not explain the broader spread of larval samples in the MDS space.

**Figure 5.**
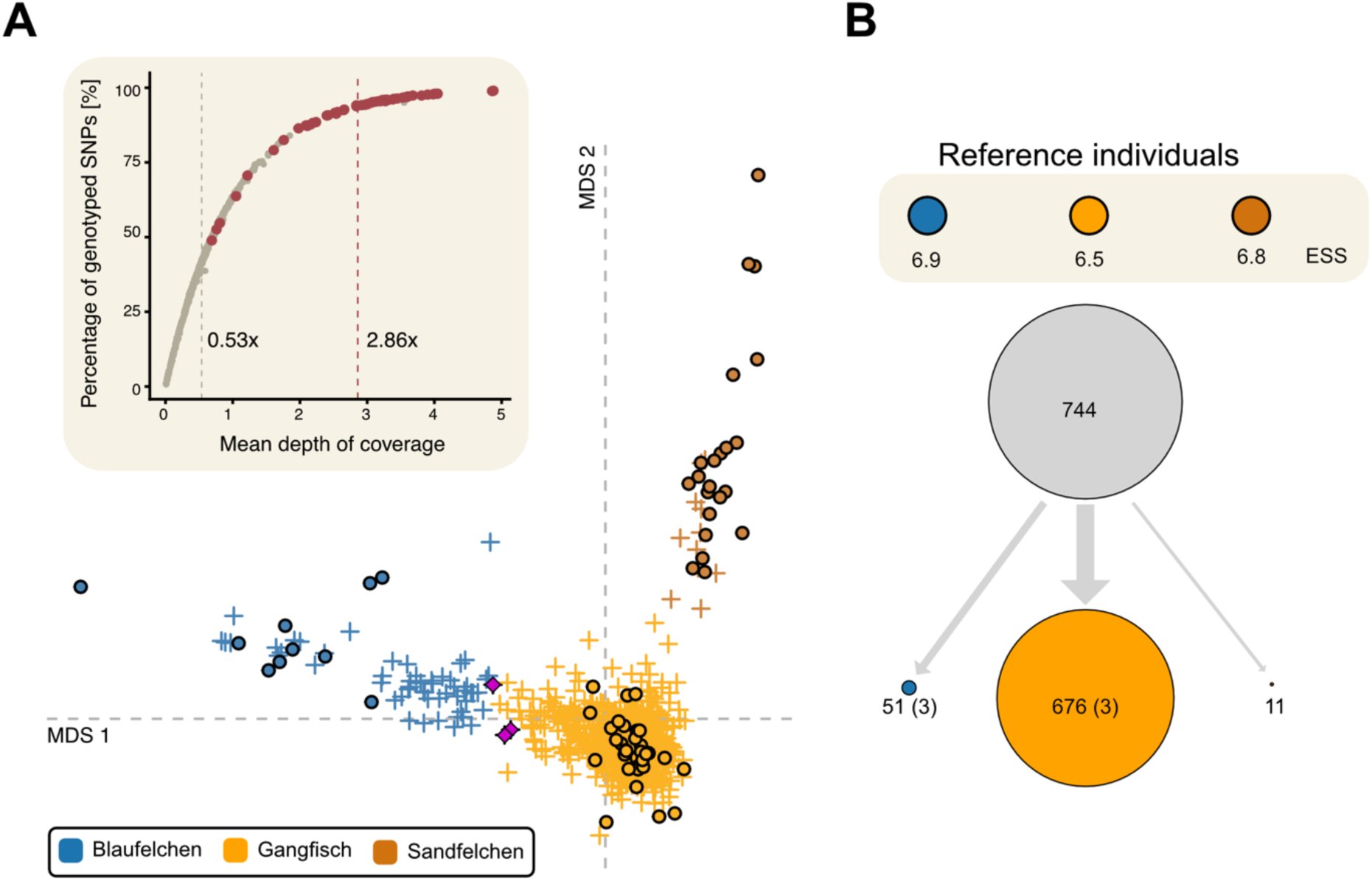
Larval assignment. (**A**) MDS plot based on filtered SNPs with larvae (crosses) coloured by species assignment. Reference individuals are shown as dots. Three larval individuals with inconsistent assignment (consistency < 0.8) are highlighted as magenta diamonds. (**B**) Species assignment results for larvae. The upper panel shows the reference populations with their respective effective sample sizes (ESS). The large grey circle shows the total number of larvae, and the lower coloured circles show the number of larvae assigned to each species, coloured by species. Circle sizes are proportional to the number of individuals assigned. Numbers in brackets show the number of individuals with assignment consistencies below 0.8.

### Spatial distribution of species in Lake Constance

To determine how species are spatially distributed across Lake Constance at different life stages, we investigated differences in spatial and depth distribution across species and lake parts. Almost all adult Gangfisch and Blaufelchen were caught in the Upper Lake, yet one Blaufelchen and ten Gangfisch were caught in the lower lake. We couldn’t genetically distinguish Sandfelchen and Unterseefelchen, suggesting that Sandfelchen and Unterseefelchen are one population (Fig. 6A).

**Figure 6.**
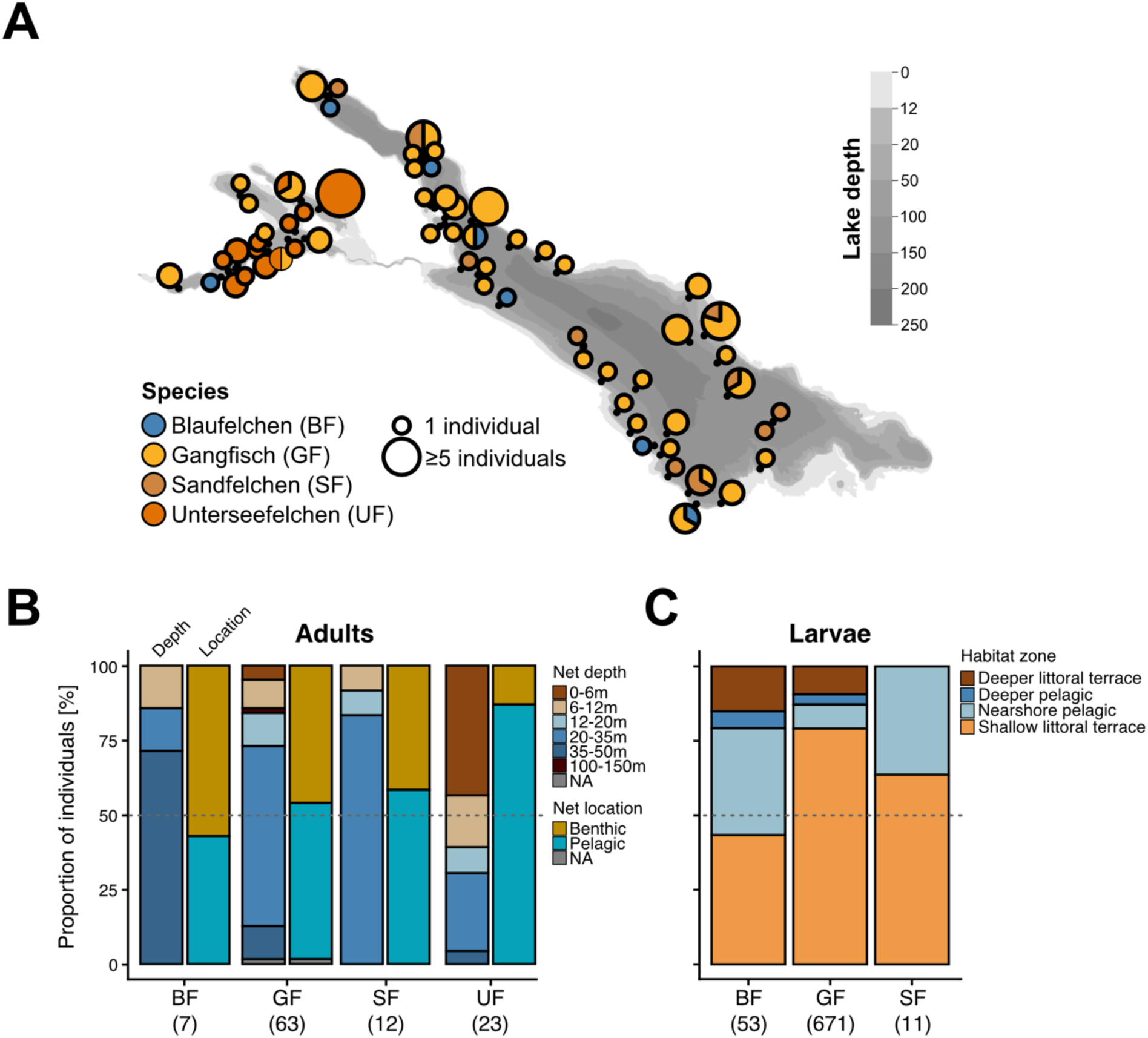
Habitat usage of adult and larval Lake Constance whitefish. (**A**) Spatial distribution of adult whitefish caught during the lake-wide fishing campaign in September 2024. Points show the location of individual nets, with the size indicating the number of individuals for each net and the colour the genetic species assignment. (**B**) The proportion of adults caught during the lake-wide fishing campaign in 2024 in nets set in different depth zones and different locations/habitats. (**C**) The proportion of larvae caught in different habitat zones by species. The numbers below the abbreviated species names in (B) and (C) show the sample size. BF = Blaufelchen, GF = Gangfisch, SF = Sandfelchen, UF = Unterseefelchen.

The depth distribution of adult whitefish caught during the lake-wide fishing campaign suggests the presence of depth preferences between species (Fig.6B). The majority of Blaufelchen were caught in deeper nets between 35 and 50m (71.4%; n=5), with smaller proportions in 20-35m (14.3%, n=1) and 6-12m (14.3%, n=1). The majority of Gangfisch (60.3%; n=38) and Sandelchen (83,3%; n=10) were caught between 20-35m, although Gangfisch seemed to be present across the entire depth distribution down to 150m (n=1 Gangfisch in a deep net). However, this could potentially be explained by the larger sample size of Gangfisch. Unterseefelchen, which are genetically Sandfelchen, were mostly caught in shallow pelagic nets below 6m (43.5%; n=10) and between 20-35m (26.1%; n=5). Similar to the adults, we detected subtle differences in habitat distribution between species as larvae. While most larvae for all species were caught in the shallow-littoral zone (43.4% to 79.1% of larvae per species; Fig.6C), a higher proportion of Blaufelchen and Sandfelchen larvae (35.8% and 36.4%, respectively) were present in the nearshore pelagic zone compared to Gangfish larvae (8.1%). While we did not detect any Sandfelchen larvae in deeper pelagic or littoral zones, we found both Blaufelchen and Gangfisch in these zones. These results suggest that Sandfelchen larvae are largely present in the shallow, nearshore zone, while both Blaufelchen and Gangfisch are present throughout all zones. However, it has to be noted that sampling bias can also impact these results, as most larvae were caught in the shallow littoral zone (72% of all larvae, n=531).

## Discussion

Here we use large-scale low- to ultra-low coverage whole-genome sequencing of adult and larval Lake Constance whitefish to characterise the contemporary genetic diversity, population trends, and species composition across entire Lake Constance at unprecedented spatial and genomic resolution. Beyond European whitefish, our results provide a general framework for genomic monitoring of fisheries-relevant populations with large population sizes and ongoing gene flow. By adapting cost-effective sequencing and analytical approaches for vertebrate-sized genomes, we demonstrate how population assignment, species composition, and genomic diversity can be assessed simultaneously, even under weak genetic differentiation. Our findings have important implications for current management practices in Lake Constance, as discussed below.

### Suitability of lcWGS for fisheries management

Genetic stock assignment remains challenging in systems with strong gene flow and limited genotyping tools (Benestan et al., 2015; Bernos et al., 2024; DeSaix et al., 2019; Theissinger et al., 2023). Reduced-representation sequencing approaches have been widely adopted but are labour-intensive, target only a small fraction of the genome, and can be difficult to integrate across studies, limiting their value for long-term monitoring (Theissinger et al., 2023). Here, we show that (ultra-)low coverage WGS enables accurate species assignment, spatial analyses, and genome-wide inference of diversity and differentiation in weakly structured whitefish populations. Despite low genome-wide differentiation, individuals could be reliably assigned to species down to sequencing depths of 0.01×, despite being genotyped at only ∼1% of total SNPs (DeSaix et al., 2023, 2024). This was enabled by a rapid and cost-effective library preparation protocol (Gaio et al., 2022), which we adapted to large genomes (*Coregonus* spp. genome: ∼2.5Gb) (De-Kayne et al., 2020). This enabled us to prepare WGS libraries for around £2.5 per library (after some minor start-up costs) in less than 1 day (for 96 samples) and sequence them to 1x coverage for approximately £7.5 per sample (sequencing cost in 2024), making it significantly cheaper and quicker than most reduced-representation sequencing approaches (e.g. RADseq) (Lou et al., 2021). Importantly, the use of whole-genome data without targeted markers facilitates seamless integration of future samples, as demonstrated by combining our dataset with previously published whitefish genomes (Frei et al., 2022). These results highlight the broad potential of lcWGS for fisheries management, including in highly connected systems, such as hybridizing species or marine organisms (Jacobs et al., 2018; Quintela et al., 2020; Seljestad et al., 2024).

### Genetic mixing of species across habitats and life-stages

Effective fisheries management, including hatchery supplementation, relies on the accurate species identification of spawning adults, often inferred from spawning location or morphology in the field. However, environmental change can erode prezygotic barriers and undermine such assumptions, as previously shown for Lake Constance whitefish <u>(Vonlanthen et al. 2012; Frei et al. 2023; Frei et al. 2022; Eckmann and Rösch 1998; Numann 1978)</u>. Targeted sampling revealed extensive mixing of Blaufelchen and Gangfisch on traditional spawning grounds, which are often used to determine the species during broodstock fisheries, which has therefore strong implications for hatchery practices that aim to create species-pure crosses (Baer et al., 2023). Notably, ∼70% of individuals caught on Blaufelchen spawning grounds were genetically Gangfisch, raising the possibility that hatchery supplementation unintentionally contributes to genetic mixing between species. Larval data further support this interpretation, showing nearly continuous genetic variation between Blaufelchen and Gangfisch, particularly at early life stages. The reduced frequency of genetically intermediate adults relative to larvae suggests possible selection against admixed individuals later in life (‘viability selection’) (Blain et al., 2024; Moser et al., 2016; Schluter et al., 2025), although targeted experimental and longitudinal studies are needed to confirm this. Viability selection against strongly admixed individuals could be one explanation for the maintenance of distinct whitefish species despite strong gene flow.

Despite historical stocking of non-endemic species, we found no evidence for genetic contributions from such introductions, consistent with previous work in Lake Constance (Baer, Schliewen, et al., 2022; Frei, Mwaiko, et al., 2023). Contemporary samples consistently cluster by species, indicating that current genetic patterns largely reflect endemic lineages.

Species misidentification is further compounded by weak morphological differentiation. Traditional phenotypic traits have long been recognised as unreliable, particularly between Blaufelchen and Gangfisch since eutrophication (Eckmann & Rösch, 1998; Hartmann & Knöpfler, 1977; Numann, 1986). Our results confirm that even advanced morphometric analyses fail to reliably distinguish these species at the individual level, and that body shape correlates with genetic admixture. While gonadal maturity and life-stage differences likely influence body shape (Helland et al. 2009), we could still distinguish Sandfelchen/Unterseefelchen from Gangfisch, suggesting that morphology reflects broad habitat use in some instances (Etheridge et al., 2012; Harrod et al., 2010). Overall, morphology and spawning habitat alone are currently unreliable for species identification in Lake Constance, underscoring the need for genomic tools and analyses.

### Skewed stock composition and habitat use

Our lake-wide species assignment of adults and larvae revealed a highly skewed stock composition, with Gangfisch comprising ∼87% of individuals. This dominance may reflect the broader ecological niche and extended spawning period of Gangfisch, which could buffer recruitment against unfavourable conditions (Frei, Reichlin, et al., 2023; Jacobs et al., 2019; Schweizer, 1894; Steinmann, 1950). In contrast, Blaufelchen - the primary commercial target - accounted for only ∼7% of sampled individuals, consistent with recent stock collapses (Haase et al., 2025). Although Sandfelchen were rare in Upper Lake Constance (∼3%), all individuals in Lower Lake Constance clustered genetically with Sandfelchen with minimal genetic differentiation, indicating that the Sandfelchen/Unterseefelchen form a single population, and should likely be managed as a single stock. Furthermore, the identification of a Sandfelchen individual in the Alpenrhein further supports regular river use by this species (Steinmann, 1950). Interestingly, Blaufelchen and Gangfisch were also detected in Lower Lake Constance during lake-wide survey in 2024, although their spawning there remains unresolved.

Habitat use seemed to differ across species and life stages, at least during the snapshot that the sampling across a few timepoints provided. Adult distributions during feeding (September 2024) were more complex than traditional classifications suggest (Steinmann, 1950), with the pelagic-planktivorous Blaufelchen frequently caught in deeper benthic nets and the littoral Sandfelchen in pelagic nets at medium depth. Gangfisch consistently occupied the broadest depth range, also below 100m, which is consistent with the hypothesised niche expansion due to introgression from the extinct Kilch (Frei et al., 2022; Frei, Reichlin, et al., 2023; Hirsch et al., 2013; Jacobs et al., 2019). However, to fully understand contemporary habitat use differences between species it will be crucial to follow genetically-confirmed individuals throughout the year, e.g. using acoustic telemetry, , as habitat use can differ throughout and across years <u>(Thomas et al. 2010)</u>.

While larvae were mostly present in the shallow nearshore habitat, larval distributions aligned more closely with expected spawning habitats <u>(Steinmann 1950)</u>, although sampling bias - particularly limited coverage of deep pelagic zones - may underestimate the abundance of Blaufelchen larvae. Larval Blaufelchen are thought to prefer deep pelagic habitats due to the lack of anti-predator mechanisms, which are less important in the deep pelagic zone (Ros et al., 2019). However, the low proportion of adult Blaufelchen in the catch still suggests a low census population size.

### Genetic diversity and genomic conservation tools

Consistent with previous work, we detected low effective population sizes and low genetic diversity (Robinson et al., 2016) across all species, with pronounced declines in Blaufelchen (Frei, Mwaiko, et al., 2023). While lower sample sizes for Blaufelchen could influence estimates, sample sizes likely reflect true census abundances (Haase et al., 2025). Given the importance of neutral and adaptive genetic diversity for population recovery and resilience (Exposito-Alonso et al., 2022; Kardos et al., 2021; Steffen et al., 2015), these findings raise concerns about the capacity of Lake Constance whitefish - particularly Blaufelchen - to respond to future environmental change.

Despite extensive genome-wide homogenisation, we identified genomic regions with elevated differentiation, including a region on chr23 overlapping the *edar* gene associated with gill raker variation (De-Kayne et al., 2022). Although allele frequencies have converged over time (Frei, Mwaiko, et al., 2023), this region remains differentiated between Sandfelchen and Gangfisch and shows signatures of selection in Blaufelchen, indicating the persistence of functional genomic variation. Many differentiated regions overlapped putative low-recombination regions rather than structural variants, suggesting that reduced recombination facilitates the persistence of adaptive divergence despite gene flow. Further work is needed to resolve the genomic architecture underlying adaptive variation in this system (De-Kayne et al., 2022; Frei et al., 2022; Frei, Reichlin, et al., 2023; Jacobs et al., 2019), which can form the basis for novel genomic monitoring tools of adaptive genetic variation and contribute to effective management decisions (Garner et al., 2016; Hendricks et al., 2018; Pearse, 2016).

## Conclusions & Limitations

By integrating lake-wide genomic data across life stages and lake basins, we show that Lake Constance whitefish stocks are highly skewed toward one species, show mixing of species on spawning grounds, and characterised by low genetic diversity and differentiation between species. Importantly, we show that the commercially important Blaufelchen makes up only a small proportion of the stock. These findings have direct implications for current hatchery practices and management strategies, which rely on accurate species identification. To fully understand the impact of mixed spawning grounds on hatchery supplementation and natural admixture, future studies should directly compare the genetic composition of hatchery and wild juveniles. While our low-coverage approach limits inference of fine-scale inbreeding patterns, it provides a powerful tool for lake-wide genomic monitoring of stock composition. Overall, our results call for a re-evaluation of current fisheries management strategies in Lake Constance.

## Supporting information

Supplementary file 1 - Protocol

## Acknowledgements

This project was funded by the Ministry of Food, Rural Area, and Consumer Protection (Baden Wurttemberg; Germany) to AB. AJ was supported by a NERC Independent Research Fellowship (NE/W008963/1).

## Author contributions

Conceptualisation: AB and AJ; Sample collection: AB and BR; Laboratory work: AJ and MC; Genomic Analysis: AJ; Phenotypic analysis: SR; Writing first draft: SR and AJ; Final draft: All authors.

## Data availability statement

All data will be made available upon acceptance of the manuscript. The raw sequencing files are accessible on ENA under the BioProject XXXXX. Additional data are available on xxxx.

