## Supplementary file 1 - Protocol for "Revisiting the genetics of Lake Constance Coregonids using lake-wide whole genome sequencing"

### Hackflex

#### Protocol to create DNA libraries for low-coverage whole genome sequencing based on the Illumina DNA prep kit.

**Reference: Gaio et al., Microbial Genomics 2022;8:000744 DOI 10.1099/mgen.0.000744**

**Modified by Arne Jacobs and Maria Capstick**

Components:

- Bead-Linked Transposomes (BLT) from illumina DNA prep kit (Stored at 4 ºC)
- Lab-made Tagmentation Buffer (LTB) (-20 ºC)
    - 20mM Tris-HCL (differs from Gaio et al)
    - 20mM MgCl
    - 20% DMF (reduced from 50%)
    - Nuclease Free Water
- Lab-made Tagment Stop Buffer (LTSB) (RT)
    - 0.2% SDS
    - Nuclease Free Water
- PrimeSTAR GXL DNA Polymerase kit (-20 ºC)
- Barcoded Index (custom design (v1) and ordered through IDT) (-20 ºC)
    - See associated file (hackflex barcodes) with 96 i5 and 96 i7 options in original paper.
- Resuspension Buffer (RSB) (4 ºC)
- Freshly prepared 80% ethanol
- Nuclease Free Water
- Ampure XP Beads (4 ºC)
- 10 µl with 10ng of DNA.

**1. Tagmentation**

- Allow BLT to equilibrate to room temp on the bench top for at least 30 minutes before use. Bring LTB to room temp.
- Add 2–10 µl DNA to each well of a 96-well PCR plate so that the total input amount is **10 ng**.
- If DNA volume < 10 µl, add nuclease-free water to the DNA samples to bring the total **volume to 10 µl**.
- Vortex BLT vigorously for to resuspend. Repeat as necessary.
- Prepare a 1:50 dilution of BLT in nuclease free water (10µl + extra per sample)
- Prepare tagmentation master mix (overage is already included):

| Reagent | Volume (µl)<br><br>(without overage) | Volume (96 samples)<br><br>(with overage) |
|---|---|---|
| 1:50 BLT | 10 | 1008 |
| LTB | 25 | 2520 |

- Vortex the tagmentation master mix thoroughly to resuspend.
- Transfer 35 µl tagmentation master mix to each well of the plate containing a sample. If prepping a large number of samples, you may divide the mix volume equally into an 8-tube strip and use a multichannel pippette to transfer.
- Pipette each sample 10 times to resuspend.
- Seal the plate, place in the preprogramed thermal cycler, and run the TAG program. Your total reaction volume will be 45µl.

**<u>TAG program:</u>**

Choose the preheat lid option and set to 100°C

- Set the reaction volume to 50 µl
- 55°C for 15 minutes
- Hold at 10°C

***2. Post-Tagmentation Cleanup***

- Add 10 µl LTSB to the tagmentation reaction.
- Slowly pipette each well 10 times to resuspend the beads.
- Seal the plate, place on the pre-programmed thermal cycler, and run the PTC program.

  **PTC program:**

  Choose the preheat lid option and set to 100°C

  Set the reaction volume to 60 µl

  37°C for 15 minutes

  Hold at 10°C

- After removing from the thermocycler, place the plate on the magnetic stand and wait until liquid is clear (~3 minutes). Keep on the magnetic stand until the appropriate step in the next section.

**3. PCR**

- Prepare PrimeStar PCR master mix, per sample (overage is already included):

| Reagent | Volume (µl) (without overage) | Volume (µl) (with overage) | Volume (µl) (96 samples; with overage) |
|---|---|---|---|
| 5x GXL Buffer | 10 | 10.5 | 1008 |
| 25mM dNTPs | 4 | 4.2 | 403.2 |
| PrimeStar GXL polymerase | 1 | 1.05 | 100.8 |
| Nuclease-free water | 20 | 21 | 2016 |

- Vortex, and then centrifuge the PCR master mix at 280 × g for 10 seconds.
- With the plate on the magnetic stand, use a 200 µl multichannel pipette to remove and discard supernatant. Foam that remains on the well walls does not adversely affect the library.
- Remove from the magnet.
- Immediately add 35 µl PrimeStar PCR master mix directly onto the beads in each sample well.
- Immediately pipette to mix until the beads are fully resuspended. Alternatively, seal the plate and use a plate shaker at 1600 rpm for 1 minute.
- Seal the sample plate and centrifuge at 280 × g for 3 seconds.

- Add the appropriate index adapters to each sample: Add 5µL each of one 10µM i7 and one 10µM i5 adapter.
- Using a pipette set to 35 µl, pipette 10 times to mix. Alternatively, seal the plate and use a plate shaker at 1600 rpm for 1 minute.
- Centrifuge at 280 × g for 30 seconds. Place on the thermal cycler and run the BLT PCR program. Your reaction volume is 45uL.

PCR program

| Temp (C) | Time (min:sec) |
|---|---|
| 68 | 3:00 |
| 98 | 3:00 |
| **98** | **0:45** |
| **62** | **0:30** |
| **68** | **2:00** |
| 68 | 1:00 |
| 10 | hold |

- The bolded steps will be repeated for the appropriate number of cycles according to the following chart.
- If doing Hackflex with 10ng of input, 12 cycles is recommended.

**4A. Library Clean-up (double-sided)**

- Allow Sample Purification Beads or Ampure XP Beads to equilibrate to room temperature on the bench for at least 30 minutes before use.
- Vortex beads before each use and frequently during use to make sure that beads are evenly distributed.
- Centrifuge plate at 280 × g for 1 minute to collect contents at the bottom of the well.
- Place the plate on the magnetic stand and wait until the liquid is clear (~5 minutes).
- Add 22.5 µl beads to each well of a new round-bottom plate
- Transfer 45 µl supernatant from each well of the PCR plate to the corresponding well of this new round-bottom plate containing beads.
    - *Due to Hackflex's lower PCR reaction volume, you may get slightly less supernatant here and in the proceeding steps. If you are not greater than 5µl off the given volumes, continue with the protocol as written.*
- Pipette each well 10 times to mix. Alternatively, seal the plate and use a plate shaker at 1600 rpm for 1 minute.
- Incubate at room temperature for 5 minutes.
- Place on the magnetic stand and wait until the liquid is clear (~5 minutes).
- During incubation, thoroughly vortex the beads, and then add 6 µl to each well of a new round-bottom plate.
- Transfer 60 µl supernatant from each well of the first plate into the corresponding well of the second plate (containing 6 µl beads).
- Pipette each well in the second plate 10 times to mix. Alternatively, seal the plate and use a plate shaker at 1600 rpm for 1 minute.
- Discard the first plate.
- Incubate at room temperature for 5 minutes.
- Place on a magnetic stand and wait until the liquid is clear (~5 minutes).
- Remove and discard all supernatant from each well without disturbing the beads.
- Wash 2 times as follows:
    - Add 180 µl fresh 80% EtOH to each well.
    - Incubate on the magnetic stand for 30 seconds.
    - Remove and discard all supernatant from each well.
- Using a 20 µl pipette, remove residual 80% EtOH from each well.

- Air-dry on the magnetic stand for 5 minutes.
- Remove from the magnetic stand.
- Add 32 µl RSB to each well.
- Pipette to resuspend.
- Incubate at room temperature for 2 minutes.
- Place on a magnetic stand and wait until the liquid is clear (~2 minutes).
- Transfer 30 µl supernatant to a new plate.

**5. Library QC**

- All samples: Qubit HS dsDNA assay
- Tapestation D1000/D5000: for subset of samples
