## Supplementary Material for "Revisiting the genetics of Lake Constance Coregonids using lake-wide whole genome sequencing"

#### Supplementary Methods:

##### *Phenotypic analysis*

To compare overall body shape among the different genetically identified whitefish species, we selected 17 homologous points (Fig. SX, Table SX) that could be clearly identified on the body of all specimens as landmarks and superimposed them on digital images using *R* 4.5.2 (R Core Team 2025) and the function *digitize2d* from the *geomorph* package (V4.0.10, Baken et al. 2021, Adams et al. 2025). The *geomorph* package was also used for the subsequent statistical analyses. Generalized Procrustes Analysis (Gower, 1975) was performed using the *gpagen* function to remove size, position and orientation differences. Whitefish with abnormal body shape (e.g. due to clear body arching) were identified using the *plotOutliers* function, which detects individuals that deviate substantially from the mean shape. Conspicuous specimens were manually checked and, if necessary, excluded from the analysis. For the final statistical analysis, we considered 158 individuals (Table 1). A model was developed, considering allometric effects (size) and species (genetic assignment): coords ~ log(size) * species. Permutation-based Procrustes ANOVA using residual randomization (functions: *procD.lm* and *anova*, permutations = 9999; estimation method: ordinary least squares; Goodall, 1991) was used to test differences in shape between species. As size had a significant effect on body shape, we used a second permutation-based Procrustes

ANOVA (permutations = 9999; estimation method: ordinary least squares) to test whether allometric effects were common among all fish or unique to a species. Since the allometric effects were different for certain species, a size correction was not performed. We then used the pairwise function of the *R* package *RRPP* (V2.1.2, Collier & Adams, 2018) as a post hoc test to compare whitefish species. In the analysis, the previously developed model was used with permutations = 9999, test type = distance between vectors (dist) and confidence = 0.95. To account for allometric effects, the null model: coords ~ log(size) was used in the pairwise analysis. The obtained p values were adjusted using the Bonferroni-Holm method (Holm, 1979). To illustrate the differences in shape between the whitefish species examined, we performed a Principal Component Analysis (PCA) using the *R* package *geomorph*. The differences in shape between the two extremes along the PC axes were visualized using wireframe models. Finally, a canonical variate analysis (CVA; Campbell & Atchley, 1981) was used to examine the accuracy of species classification based on body shape, using the *prep.lda* function of the *R* package *RRPP* and the *CVA* function of the *R* package *Morpho* (V2.13, Schlager, 2013). For the analysis, n – 1 PCs (n = number of individuals in the smallest population) were taken into account and a jackknife cross-validation (Fan & Wang, 1996) was performed to validate the group assignments.

Phenotypic differences between whitefish species were further investigated using 15 linear measurements (Fig. SX, Table SX). The measurements were taken from digital images of 158 individuals (Table 1) using *ImageJ* (V1.53c, Schindelin et al. 2012). Size correction was conducted based on the standard length in *R* using the *GroupStruct* package (V0.1.0, Chan & Grismer). The size correction method uses an allometric growth equation according to Thorpe (1975), whereby the slope and the mean body length was calculated jointly for all individuals examined. The analyses described below were repeated, with the slope being calculated separately for each whitefish species and with both the slope and the average body length being calculated separately for each species. To identify linear measurements with a high correlation, a correlation matrix (correlation coefficient: Spearman rank correlation) was calculated based on the size-corrected measurements using the *cor_mat* function of the *rstatix* package (V0.7.3, Kassambara 2023). Measurements exhibiting a correlation coefficient >0.9 were excluded from the dataset. Furthermore, we used Recursive Feature Elimination (RFE) to identify linear measurements that contribute most to the differentiation of the whitefish species. For this, the *rfe* function of the *R* package *caret* (V7.0.1, Kuhn 2008) was used with the following settings: model function = Random Forest, method = repeated cross-validation, number of folds = 10, number of repeats = 10. Measurements selected in the model with the highest accuracy were utilized in the subsequent analyses. For these, we used the *R* package *MorphoTools2* (V1.0.2.1, Šlenker et al. 2022). The selected linear measurements were used in a PCA (function: *pca.calc*) and the first two PCs, which explained the greatest variance in shape, were graphically represented in scatter plots. To investigate the success of species classification based on the selected linear measurements, we performed a K nearest neighbor classificatory discriminant analysis (distance: Euclidean). The optimal number of neighbors was estimated empirically using the function *knn.select*. Cross validation (mode: individual) was then performed using the function *classif.knn* of the *MorphoTools2* package.

***Whole genome sequencing***

DNA was extracted from fin clips for larval and adult samples using magnetic beads following the protocol by Kučka and Chan (dx.doi.org/10.17504/protocols.io.b46bqzan) with minor modifications. Fin clips were digested overnight at 56°C in a shaking oven using 10µl of Proteinase K and 90µl of PureLink Digestion buffer. We omitted the RNAse treatment step due to the absence of intact RNA in samples and reduced the centrifugation to pellet undigested material to 1 min due to the absence of undigested material in the sample. Furthermore, washing steps of beads were performed with just 50% and 80% ethanol and DNA was eluted into 60µl of elution buffer. Following DNA extraction, DNA concentration was quantified using a Qubit Flex with the 1X BR dsDNA assay and quality was determined using agarose gels.

Whole genome sequencing (WGS) libraries for low-coverage WGS (lcWGS) were prepared using a modified HackFlex protocol (Gaio et al. 2022) (see Supplementary Material for protocol). In brief, we used 10ng of DNA as input (1ng/µl) and diluted the illumina DNA prep BLT beads 1:50 in nuclease-free water. However, we reduced the amount of DMF in the tagmentation buffer to 20% (compared to 50% in the original protocol) to reduce the amount of this highly toxic chemical without impacting library quality. The tagmentation was performed as described in the original protocol and we skipped the clean-up step prior to the indexing PCR. The indexing PCR was performed for 12 cycles using custom i5 and i7 barcodes (see original paper). In contrast to the original protocol, we performed a double-sided bead clean up on the final libraries using 0.5X and 0.1X Ampure XP Beads. The final libraries were quantified using the Qubit Flex with the HS dsDNA assay and a subset of libraries were run on the Tapestation using the D1000 or D5000 tapes to check the fragment size distribution, with an expected peak around 400-600bp. We pooled libraries at equal molarities, using the average fragment length distribution for molarity calculations. We sequenced up to 288 pooled libraries using 150PE reads on NovaSeq X Plus 10B lanes at Novogene UK to an average of 1.6 Gb per individual. Some individuals were sequenced on an additional NovaSeq X Plus 25B lane to increase the coverage.

Furthermore, we downloaded public whole genome sequencing data for adult individuals with confirmed species assignment from NCBI SRA for Gangfish (C. macrophthalmus), Sandfelchen (C. arenicolus), and Blaufelchen (C. wartmanni) from Upper Lake Constance (Frei et al. 2023, 2022) (Table S2).

***Bioinformatic processing and variant identification***

Raw sequencing reads were processed following best practices for lcWGS data (Lou et al. 2021) by first trimming low quality bases, adapters and poly-G tails using fastp using the -l 30 --trim_poly_g --cut_right --detect_adapter_for_pe settings (Chen et al., 2018). We mapped the processed reads to the European whitefish ('Balchen') reference genome (AWGv2; (De-Kayne & Zoller, 2020)) using *bwa mem* (Li & Durbin, 2009), merged replicated samples using the merge command in *sambamba* v.0.8.2 (Tarasov et al., 2015), removed duplicated reads with *sambamba* using *--remove-duplicates*, clipped overlapping reads using the *bamUtils clipOverlap* command (https://github.com/statgen/bamUtil), and indexed the bam files using *sambamba index*. Finally, we performed indel realignment per sample using GATK3.8 with the RealignerTargetCreator and IndelRealigner commands. We estimated the depth of coverage for each bam file using *mosdepth* v.0.3.11 with the -n --fast-mode --by 500 --mapq 30 flags (Pedersen & Quinlan, 2018).

Due to the low coverage of the sequencing data, we performed genotype likelihood-based (GL) analyses with *ANGSD* v0.938 (Korneliussen et al., 2014). We created two different SNP datasets to address distinct questions. Both datasets used the same filtering strategies. For the first list, we called SNPs across all 250 adult samples ('adult SNP dataset'). We identified variant sites across all adult samples using ANGSD with the samtools genotype likelihood model (-GL 1) with a minimum SNP value of 1e-6 (-SNP_pval 1e-6), determining major/minor alleles based on allele frequencies (-doMajorMinor 1), removing sites below and above a minimum and maximum total sequencing depth of 250 (minimum number of individuals used to call SNPs x mean sequencing depth) and 1300 (2 x total number of individuals x mean sequencing depth), respectively, across all individuals, and removed sites with missing data in more than 20% of individuals (-minInd 200), minimum base quality below 30 (-minQ 30) and a minimum individual depth of 1 (-setMinDepthInd 1). Furthermore, we removed multi-mapping reads (-uniqueOnly 1), unmapped or duplicated reads (-remove_bads 1) and those without a matching paired read (-

only_proper_pairs 1). We also adjusted the mapping quality for excessive mismatches (-C 50). We only kept SNPs with a minor allele frequency of 5% (-minMaf 0.05). We then removed SNPs falling within potentially problematic genomic regions that can impact analyses accuracy. First, we removed deviant SNPs, meaning those with excess heterozygosity potentially due to collapsed paralogs, using the ngsParalog calcLR command based on samtools mpileup files (Dallaire et al. 2023). Second, we excluded all sites that fall within likely collapsed regions within the genome assembly (Frei et al. 2023). Third, we removed those sites that fall within regions without uniuqe mappability (Wang et al. 2024), as inferred using a kmer approach in genmap (150-mers) (Pockrandt et al. 2020).

For the second list, we created an additional SNP dataset for a subset of adult reference samples (published data and individuals from targeted sampling) that was used for species assignments of larvae ('larval SNP dataset'). Some adult individuals showed wrong species assignment in the field and species assignments were corrected based on the genomic data (see results below). We identified variant sites across adult reference samples (Table S2) using ANGSD with the same filtering strategy as above but adjusted minimum depth and missingness filters to a minimum sequencing depth to 300, maximum depth of 600, and missingness of below 20% of individuals (-minInd 66). Since larvae had ultra-low sequencing coverage, we did not use them for SNP identification, but instead inferred genotype likelihoods for larvae based on the filtered 'larval SNP dataset' with the -sites function, exporting GLs in beagle format (-doGlf 2). We did not apply any other filters at this step, such as depth filters, due to the very low coverage (DeSaix et al. 2024).

**Selective sweeps**

The PBE is an extension of the population branch statistic (PBS), which aims to identify selective sweep signatures on individual genomic regions by comparing the focal branch length of a locus within a population compared to two outgroup populations (Shpak et al., 2025). PBE extends this approach and estimates selective sweeps by testing if the observed PBS value exceeds the expected value. PBE was estimated as: $PBE_A = PBS_A - PBS_{A\text{-}exp} = PBS_A - [(T_{BC} \times med(PBS_A)) / med(T_{BC})]$, with $med(PBS_A)$ and $med(T_{BC})$ being the median values of their respective statistics across all loci and $T_{BC}$ being the linearised Fst between population B and C at this locus.

***Structural variant identification***

To identify putative low recombination regions and structural variants across the whitefish genome that might shape the landscape of genetic diversity and differentiation, we employed a local PCA approach in LOSTRUCT (Li & Ralph, 2019). To accommodate genotype likelihoods (GLs), we implemented an approach that performs PCA on fixed windows across the genome using PCAngsd (Zhou et al., 2024). In brief, the beagle file was divided into 250 SNP windows and PCAs performed per window using PCAngsd. These PCAs were used as input for LOSTRUCT to estimate distances between windows and perform multi-dimensional scaling, and outlier windows were identified as those with absolute z-transformed MDS values above 4 (≥ 4 standard deviations above the mean). For outlier regions that were over 1Mb in size after merging adjacent outlier windows (23 regions in total), we performed PCA using PCAngsd to further test for genetic clustering patterns that are indicative of structural variants, such as inversions. Inversions usually create a PCA pattern with three distinct clusters, corresponding to the three distinct karyotypes of inversions.

### Supplementary Files

**Supplementary file 1:** Modified Hackflex protocol used for the DNA library preparation.

### Supplementary Tables

**Table S1.** Overview of the targeted fishing campaign, targeting adult individuals of various whitefish species occurring in Lake Constance.

| **Location** | **Sampling date** | **n samples (genetics)** | **n samples (morpho-metrics)** | **Coords (Easting)** | **Coords (Northing)** |
|---|---|---|---|---|---|
| Upper Lake (pelagic zone)[a] | 08.12.2021 | 17 | 17 | 537779 | 5266907 |
| Upper Lake (littoral zone)[a] | 15.12.2021 | 20 | 20 | 5407781 | 5270998 |
| Upper Lake (littoral zone)[b] | 10.03.2021-1.12.2023 | 38 | 29 | 540227 | 5271406 |
| Lower Lake (“Gnadensee”) | 25.05. / 27.06.2023 | 15 | 14 | 54373 | 5284076 |
| Lower Lake (“Zellersee”) | 27.06.2023 | 15 | 15 | 499819 | 5284529 |
| Lower Lake (“Rheinsee east” | 21.07.2023 | 14 | 13 | 501344 | 5280826 |
| Lower Lake (“Rheinsee west” | 16.08.2023 | 14 | 14 | 491756 | 5277506 |
| Rhine River (“Alpenrhein”)[c] | Oct 2022 / 1988-1990 | 1 | 0 | 545069 | 5243223 |

[a] Individuals were caught during annual spawning fisheries. [b] Several fishing events took place during the period listed in order to specifically catch “Sandfelchen”. [c] One individual was caught by a local angler in 2022. The remaining individuals were caught during routine fishing events in the period listed and only scale samples were available.

**Table S2**. Overview of whitefish caught in Lake Constance in September 2024 as part of a lake-wide fishing campaign.

| Location | Sampling date | Depth [m] | Coords (Easting) | Coords (Northing) |
|---|---|---|---|---|
| Lower Lake | 01.09.2024 | 27 | 492616 | 5277251 |
| Lower Lake | 01.09.2024 | 27 | 492616 | 5277251 |
| Lower Lake | 01.09.2024 | 8 | 503071 | 5284730.5 |
| Lower Lake | 01.09.2024 | 8 | 503071 | 5284730.5 |
| Lower Lake | 01.09.2024 | 8 | 503071 | 5284730.5 |
| Lower Lake | 01.09.2024 | 37 | 496225.5 | 5278224.5 |
| Lower Lake | 01.09.2024 | 18 | 497874.5 | 5285214.5 |
| Lower Lake | 01.09.2024 | 12 | 497867 | 5285160 |
| Lower Lake | 02.09.2024 | 18 | 503359 | 5281497 |
| Lower Lake | 02.09.2024 | 18 | 503359 | 5281497 |
| Lower Lake | 02.09.2024 | 22 | 500290.5 | 5281050 |
| Lower Lake | 03.09.2024 | 12 | 496869 | 5278684.5 |
| Lower Lake | 03.09.2024 | 12 | 496869 | 5278684.5 |
| Lower Lake | 03.09.2024 | 30 | 500278.5 | 5280397.5 |
| Lower Lake | 03.09.2024 | 30 | 500278.5 | 5280397.5 |
| Lower Lake | 03.09.2024 | 18 | 500271 | 5280347.5 |
| Lower Lake | 03.09.2024 | 30 | 497645.5 | 5279186 |
| Lower Lake | 03.09.2024 | 30 | 497645.5 | 5279186 |
| Lower Lake | 01.09.2024 | 41 | 497476.5 | 5278938.5 |
| Lower Lake | 01.09.2024 | 12 | 503055.5 | 5284677.5 |
| Lower Lake | 02.09.2024 | 8 | 504569.5 | 5284009.5 |
| Lower Lake | 02.09.2024 | 8 | 504569.5 | 5284009.5 |
| Lower Lake | 02.09.2024 | 8 | 504569.5 | 5284009.5 |
| Lower Lake | 02.09.2024 | 8 | 504569.5 | 5284009.5 |
| Lower Lake | 02.09.2024 | 8 | 504569.5 | 5284009.5 |
| Lower Lake | 02.09.2024 | 8 | 504569.5 | 5284009.5 |
| Lower Lake | 02.09.2024 | 8 | 504569.5 | 5284009.5 |
| Lower Lake | 02.09.2024 | 8 | 504569.5 | 5284009.5 |
| Lower Lake | 03.09.2024 | 24 | 501937.5 | 5281442.5 |
| Lower Lake | 03.09.2024 | 20 | 502556 | 5282096.5 |
| Lower Lake | 03.09.2024 | 8 | 500251 | 5280243 |
| Lower Lake | 03.09.2024 | 18 | 496862.5 | 5278736.5 |
| Lower Lake | 03.09.2024 | 12 | 500261 | 5280295.5 |
| Lower Lake | 03.09.2024 | 12 | 500261 | 5280295.5 |
| Upper Lake | 08.09.2024 | 12 | 527154.5 | 5272370.5 |
| Upper Lake | 08.09.2024 | 46 | 524862.5 | 5278581.5 |
| Upper Lake | 08.09.2024 | 26 | 528667.5 | 5269312 |
| Upper Lake | 08.09.2024 | 46 | 527145.5 | 5272213.5 |
| Upper Lake | 08.09.2024 | 45 | 523230.5 | 5279782 |
| Upper Lake | 09.09.2024 | 30 | 517629 | 5283002.5 |
| Upper Lake | 09.09.2024 | 30 | 517629 | 5283002.5 |

| | | | | |
|---|---|---|---|---|
| Upper Lake | 09.09.2024 | 30 | 517629 | 5283002.5 |
| Upper Lake | 09.09.2024 | 30 | 517629 | 5283002.5 |
| Upper Lake | 09.09.2024 | 30 | 517629 | 5283002.5 |
| Upper Lake | 09.09.2024 | 30 | 518225.5 | 5278402 |
| Upper Lake | 09.09.2024 | 12 | 520806.5 | 5280765 |
| Upper Lake | 10.09.2024 | 25 | 514445.5 | 5284097 |
| Upper Lake | 10.09.2024 | 25 | 514445.5 | 5284097 |
| Upper Lake | 10.09.2024 | 12 | 505265 | 5293644 |
| Upper Lake | 10.09.2024 | 12 | 505265 | 5293644 |
| Upper Lake | 10.09.2024 | 12 | 505265 | 5293644 |
| Upper Lake | 10.09.2024 | 18 | 513521 | 5288315.5 |
| Upper Lake | 10.09.2024 | 30 | 513481 | 5288246.5 |
| Upper Lake | 10.09.2024 | 30 | 513481 | 5288246.5 |
| Upper Lake | 10.09.2024 | 30 | 513481 | 5288246.5 |
| Upper Lake | 09.09.2024 | 33 | 514886 | 5282529.5 |
| Upper Lake | 09.09.2024 | 51 | 515304.5 | 5282718.5 |
| Upper Lake | 09.09.2024 | 34 | 516693 | 5278893 |
| Upper Lake | 09.09.2024 | 46 | 517576.5 | 5282969.5 |
| Upper Lake | 09.09.2024 | 46 | 517576.5 | 5282969.5 |
| Upper Lake | 09.09.2024 | 43 | 519932 | 5275787 |
| Upper Lake | 09.09.2024 | 18 | 518147 | 5278410.5 |
| Upper Lake | 10.09.2024 | 30 | 513481 | 5288246.5 |
| Upper Lake | 10.09.2024 | 26 | 514092.5 | 5284335.5 |
| Upper Lake | 10.09.2024 | 29 | 515334 | 5283231.5 |
| Upper Lake | 10.09.2024 | 29 | 515334 | 5283231.5 |
| Upper Lake | 10.09.2024 | 46 | 505485.5 | 5293650 |
| Upper Lake | 10.09.2024 | 30 | 505411.5 | 5293655.5 |
| Upper Lake | 10.09.2024 | 46 | 513445.5 | 5288190.5 |
| Upper Lake | 10.09.2024 | 8 | 513584 | 5288438.5 |
| Upper Lake | 10.09.2024 | 12 | 513556 | 5288374 |
| Upper Lake | 15.09.2024 | 30 | 536207 | 5272662.5 |
| Upper Lake | 15.09.2024 | 30 | 536207 | 5272662.5 |
| Upper Lake | 15.09.2024 | 30 | 536207 | 5272662.5 |
| Upper Lake | 15.09.2024 | 30 | 529993 | 5266673.5 |
| Upper Lake | 16.09.2024 | 30 | 539406.5 | 5268071 |
| Upper Lake | 16.09.2024 | 30 | 539406.5 | 5268071 |
| Upper Lake | 16.09.2024 | 30 | 539406.5 | 5268071 |
| Upper Lake | 16.09.2024 | 30 | 534251.5 | 5264855 |
| Upper Lake | 15.09.2024 | 18 | 531590 | 5268648.5 |
| Upper Lake | 15.09.2024 | 22 | 533690 | 5263037 |
| Upper Lake | 15.09.2024 | 20 | 531129.5 | 5264912.5 |
| Upper Lake | 15.09.2024 | 48 | 533183 | 5263849.5 |
| Upper Lake | 16.09.2024 | 28 | 536140.5 | 5276449.5 |
| Upper Lake | 16.09.2024 | 28 | 536140.5 | 5276449.5 |
| Upper Lake | 16.09.2024 | 137 | 538553.5 | 5270881.5 |

| | | | | |
|---|---|---|---|---|
| Upper Lake | 16.09.2024 | 45 | 542215.5 | 5261866 |
| Upper Lake | 17.09.2024 | 30 | 543308.5 | 5265823 |
| Upper Lake | 17.09.2024 | 27 | 536772 | 5258851.5 |
| Upper Lake | 17.09.2024 | 27 | 536772 | 5258851.5 |
| Upper Lake | 17.09.2024 | 27 | 536772 | 5258851.5 |
| Upper Lake | 17.09.2024 | 29 | 536086 | 5259789.5 |
| Upper Lake | 17.09.2024 | 29 | 536086 | 5259789.5 |
| Upper Lake | 17.09.2024 | 29 | 536086 | 5259789.5 |
| Upper Lake | 16.09.2024 | 30 | 534251.5 | 5264855 |
| Upper Lake | 17.09.2024 | 18 | 543320.5 | 5265896.5 |
| Upper Lake | 17.09.2024 | 28 | 538878 | 5258909 |
| Upper Lake | 17.09.2024 | 28 | 538878 | 5258909 |
| Upper Lake | 17.09.2024 | 28 | 537433.5 | 5273229 |
| Upper Lake | 17.09.2024 | 28 | 537433.5 | 5273229 |
| Upper Lake | 17.09.2024 | 28 | 537433.5 | 5273229 |
| Upper Lake | 17.09.2024 | 28 | 537433.5 | 5273229 |
| Upper Lake | 17.09.2024 | 28 | 537433.5 | 5273229 |
| Upper Lake | 17.09.2024 | 31 | 534363 | 5261196 |

**Table S3.** Overview of larval whitefish caught in Lake Constance in 2021 and 2022 for species classification.

| Location | Sampling year | Method | Environment | n | Corrds (easting) | Coords (northing) |
|---|---|---|---|---|---|---|
| AT2 | 2021 | Light trap | Deeper pelagic | 1 | 545909.7 | 5262165.7 |
| AT3 | 2021 | Light trap | Deeper pelagic | 3 | 545142.0 | 5263775.0 |
| AT3 | 2021 | Light trap | Nearshore pelagic | 3 | 546650.9 | 5265571.8 |
| AT4 | 2021 | Light trap | Nearshore pelagic | 15 | 546050.3 | 5262498.0 |
| AT7 | 2021 | Light trap | Nearshore pelagic | 2 | 546240.7 | 5262718.0 |
| BW1 | 2021 | Light trap | Deeper pelagic | 12 | 541493.3 | 5268564.2 |
| BW10 | 2021 | Light trap | Deeper pelagic | 1 | 533264.2 | 5271691.3 |
| BW3 | 2021 | Light trap | Deeper pelagic | 3 | 533430.0 | 5271908.4 |
| BW5 | 2021 | Light trap | Nearshore pelagic | 1 | 541019.1 | 5270629.3 |
| BW5 | 2021 | Light trap | Nearshore pelagic | 1 | 539579.0 | 5271703.2 |
| BW7 | 2021 | Light trap | Nearshore pelagic | 13 | 544339.4 | 5269703.5 |
| BW8 | 2021 | Light trap | Nearshore pelagic | 7 | 539009.0 | 5272696.4 |
| BW9 | 2021 | Light trap | Nearshore pelagic | 4 | 539446.8 | 5271576.7 |
| BY1 | 2021 | Light trap | Nearshore pelagic | 1 | 550732.6 | 5264391.4 |
| BY2 | 2021 | Light trap | Nearshore pelagic | 1 | 550699.0 | 5265361.7 |
| BY3 | 2021 | Light trap | Deeper pelagic | 1 | 547330.5 | 5264348.0 |
| DLRG | 2021 | Hand net | Shallow littoral terrace, Deeper littoral terrace | 54 | 539400.1 | 5272916.9 |
| ERNW | 2021 | Hand net | Shallow littoral terrace | 58 | 537603.1 | 5277013.2 |
| ERSE | 2021 | Hand net | Shallow littoral terrace | 25 | 538857.2 | 5274560.5 |
| LGN | 2021 | Hand net | Shallow littoral terrace | 2 | 540243.1 | 5271779.0 |
| MB | 2021 | Hand net | Shallow littoral terrace | 4 | 531988.8 | 5278708.0 |
| RS | 2021 | Hand net | Shallow littoral terrace | 47 | 547093.6 | 5260472.7 |
| SG1 | 2021 | Light trap | Nearshore pelagic | 3 | 535754.0 | 5260186.9 |
| SG2 | 2021 | Light trap | Nearshore pelagic | 1 | 539231.7 | 5259025.5 |
| SG2 | 2021 | Light trap | Deeper pelagic | 1 | 534903.8 | 5264655.1 |
| SG3 | 2021 | Light trap | Nearshore pelagic | 6 | 539792.7 | 5260926.1 |
| TG2, TG3 | 2021 | Light trap | Nearshore pelagic | 11 | 528152.7 | 5269365.4 |
| TG7 | 2021 | Light trap | Nearshore pelagic | 2 | 528184.0 | 5269379.8 |
| TG9 | 2021 | Light trap | Nearshore pelagic | 1 | 528241.2 | 5269390.7 |
| WB | 2021 | Hand net | Shallow littoral terrace | 132 | 547418.6 | 5268609.4 |
| AT01 | 2022 | Light trap | Deeper pelagic | 1 | 545056.1 | 5263832.9 |
| AT02, AT03 | 2022 | Light trap | Deeper pelagic | 2 | 545142.0 | 5263775.0 |
| BW5L, BW8L | 2022 | Light trap | Deeper littoral terrace | 4 | 539107.7 | 5272583.6 |
| BW6L, BW9L | 2022 | Light trap | Deeper littoral terrace | 10 | 539579.0 | 5271703.2 |
| BY1L, BY2L | 2022 | Light trap | Deeper littoral terrace | 3 | 546751.0 | 5268413.1 |
| BY1P | 2022 | Light trap | Deeper pelagic | 1 | 546609.3 | 5264960.9 |

| | | | | | | |
|---|---|---|---|---|---|---|
| BY2P | 2022 | Light trap | Nearshore pelagic | 1 | 549219.2 | 5265023.1 |
| BY2P | 2022 | Light trap | Nearshore pelagic | 1 | 549935.2 | 5265062.5 |
| BY3L | 2022 | Light trap | Deeper littoral terrace | 2 | 548086.7 | 5267734.8 |
| BY6L | 2022 | Light trap | Deeper littoral terrace | 1 | 547994.6 | 5267815.5 |
| DLRG | 2022 | Hand net | Shallow littoral terrace, Deeper littoral terrace | 111 | 539400.1 | 5272916.9 |
| ERNW | 2022 | Hand net | Shallow littoral terrace | 34 | 537603.1 | 5277013.2 |
| ERSE | 2022 | Hand net | Shallow littoral terrace | 21 | 538857.2 | 5274560.5 |
| IMST | 2022 | Hand net | Shallow littoral terrace | 1 | 527477.2 | 5278798.7 |
| RS | 2022 | Hand net | Shallow littoral terrace | 35 | 547093.6 | 5260472.7 |
| SG1L, SG2L | 2022 | Light trap | Deeper littoral terrace | 4 | 533818.9 | 5261142.3 |
| SG2L | 2022 | Light trap | Deeper littoral terrace | 8 | 533667.1 | 5261567.5 |
| TG10, TG6, TG8 | 2022 | Light trap | Nearshore pelagic | 3 | 528184.0 | 5269379.8 |
| TG3 | 2022 | Light trap | Deeper littoral terrace | 1 | 528225.7 | 5269381.0 |
| V01 | 2022 | Light trap | Deeper littoral terrace | 6 | 546264.8 | 5262289.8 |
| V03 | 2022 | Light trap | Deeper littoral terrace | 1 | 546280.9 | 5262235.3 |
| V04 | 2022 | Light trap | Deeper littoral terrace | 1 | 546056.9 | 5262062.1 |
| V05 | 2022 | Light trap | Deeper littoral terrace | 1 | 546520.3 | 5262263.1 |
| WB | 2022 | Hand net | Shallow littoral terrace | 76 | 547418.6 | 5268609.4 |

**Table S4 –** Overview of public whitefish whole genome sequencing data used in this manuscript. Data was downloaded from the ENA SRA. All data are from Frei et al. (2022) in Nature Ecology and Evolution.

| Species | Sampling Year | Lab ID | ENA accession | Coverage |
|---|---|---|---|---|
| C. wartmanni | 2015 | 121 | ERS6670439 | 59.4 |
| C. wartmanni | 2015 | 122 | ERS6670440 | 47.5 |
| C. wartmanni | 2015 | 123 | ERS6670441 | 58.9 |
| C. arenicolus | 2015 | 127 | ERS6670449 | 57.9 |
| C. arenicolus | 2015 | 128 | ERS6670450 | 46.4 |
| C. wartmanni | 2015 | 131 | ERS6670442 | 60.2 |
| C. macrophthalmus | 2015 | 132 | ERS6670445 | 40.5 |
| C. macrophthalmus | 2019 | 212608 | ERS12047271 | 8.2 |
| C. macrophthalmus | 2019 | 212623 | ERS12047279 | 8 |
| C. arenicolus | 2019 | 212633 | ERS12047285 | 10.3 |
| C. macrophthalmus | 2019 | 212663 | ERS12047313 | 4.3 |

**Table S5 – Landmarks** Description of the homologous landmarks used in the phenotypic analysis to discriminate between the identified whitefish species. See also Figure S1A.

| Landmark | Description |
|---|---|
| 1 | Tip of the snout |
| 2 | Most posterior part of the frontal head bone |
| 3 | Anterior insertion point of the dorsal fin |
| 4 | Posterior end of the dorsal fin (not necessarily posterior insertion point of the dorsal fin) |
| 5 | Anterior insertion point of the adipose fin |
| 6 | Posterior end of the adipose fin (not necessarily posterior insertion point of the adipose fin) |
| 7 | Dorsal insertion point of the caudal fin |
| 8 | Center point of the posterior end of the hypural plate |
| 9 | Ventral insertion point of the caudal fin |
| 10 | Posterior insertion point of the anal fin |
| 11 | Anterior insertion point of the anal fin |
| 12 | Anterior insertion point of the pelvic fin |
| 13 | Dorsal insertion point of the pectoral fin |
| 14 | Most ventral point of the body perpendicular to the dorsal insertion of the pectoral fin |
| 15 | Most ventral point of the posterior operculum margin |
| 16 | Center of the eye |
| 17 | Most dorsal point of the posterior operculum margin |

**Table S6 – Linear measurements** Description of the linear measurements used in the phenotypic analysis to discriminate between the identified whitefish species. See also Figure S1B.

| Linear measurement | Acronym | Description |
|---|---|---|
| Predorsal length | PreD | Length from the tip of the snout to the anterior insertion point of the dorsal fin |
| Postdorsal length | PostD | Length from the anterior insertion point of the dorsal fin to the posterior end of the hypural plate |
| Dorsal to adipose length | DtA | Length from the anterior insertion point of the dorsal fin to the anterior insertion point of the adipose fin |
| Caudal peduncle depth | CD | Vertical distance between the dorsal and ventral margins of the caudal peduncle at its narrowest part |
| Preanal length | PreA | Length from the tip of the snout to the anterior insertion point of the anal fin |
| Postanal length | PostA | Length from the anterior insertion point of the anal fin to the posterior end of the hypural plate |
| Anal fin base | AFB | Length between the insertions of the anal fin |
| Prepelvic length | PrePl | Length from the tip of the snout to the anterior insertion point of the pelvic fin |
| Prepectoral length | PrePc | Length from the tip of the snout to the dorsal insertion point of the pectoral fin |
| Head length | HL | Length from the tip of the snout to the most posterior point of the operculum margin |
| Eye diameter | ED | Horizontal distance across the midline of the eye from the anterior to the posterior margin of the eye |
| Snout length | SN | Length from the tip of the snout to the anterior margin of the eye |
| Postorbital length | PostO | Length from the posterior margin of the eye to the most posterior point of the operculum |
| Body depth | BD | Vertical distance between the dorsal and ventral margins of the body from the anterior insertion point of the dorsal fin to the anterior insertion of the pelvic fin (not necessarily the greatest body depth) |
| Total length | TL | Length from the tip of the snout to the tip of the longest unbranched ray either being on the dorsal or ventral part of the caudal fin |
| Standard length | SL | Length from the tip of the snout to the posterior end of the hypural plate |

**Table S7 - Genome-wide Fst between species.** BF = Blaufelchen, GF = Gangfisch, SF = Sandfelchen; UF = Unterseefelchen.

| Fst | **BF** | **GF** | **SF** | **UF** |
|---|---|---|---|---|
| **BF** | 0 | | | |
| **GF** | 0.019 | 0 | | |
| **SF** | 0.034 | 0.014 | 0 | |
| **UF** | 0.043 | 0.021 | 0.006 | 0 |

**Table S8** – Basic measurements mean weight and total length for all whitefish species.

| Species | Mean weight ± SD (g) | Mean total length ± SD (cm) |
|---|---|---|
| Blaufelchen | 292,5 ± 100.6 | 35.0 ± 8.0 |
| Gangfisch | 220.7 ± 139.2 | 29.6 ± 5.7 |
| Sandfelchen | 1318.1 ± 697.6 | 48.4 ± 10.9 |
| Unterseefelchen | 318.3 ± 123.4 | 32.3 ± 5.9 |

**Table S9** – Phenotypic classification accuracy Overview of classification accuracy for each whitefish species based on phenotypic characteristics. For landmark data, accuracy was derived from a canonical variate analysis and jackknife cross-validation. For selected linear measurements, accuracy was derived from a K nearest neighbor classificatory discriminant analysis.

| | Species | n | Blaufelchen | Gangfisch | Sandfelchen | Untersee-felchen | Correct (%) |
|---|---|---|---|---|---|---|---|
| Landmarks | Blaufelchen | 9 | 3 | 4 | 1 | 1 | 33.3 |
| | Gangfisch | 54 | 1 | 38 | 5 | 10 | 70.4 |
| | Sandfelchen | 34 | 1 | 7 | 18 | 9 | 51.4 |
| | Unterseefelchen | 61 | 0 | 6 | 9 | 45 | 75.0 |
| | Total | 158 | 5 | 55 | 33 | 65 | 65.8 |
| Linear measurements | Blaufelchen | 9 | 0 | 8 | 0 | 1 | 0.0 |
| | Gangfisch | 54 | 0 | 37 | 9 | 8 | 68.5 |
| | Sandfelchen | 34 | 0 | 11 | 11 | 12 | 32.4 |
| | Unterseefelchen | 61 | 0 | 3 | 7 | 51 | 83.6 |
| | Total | 158 | 0 | 59 | 27 | 72 | 62.7 |

### Supplementary Figures

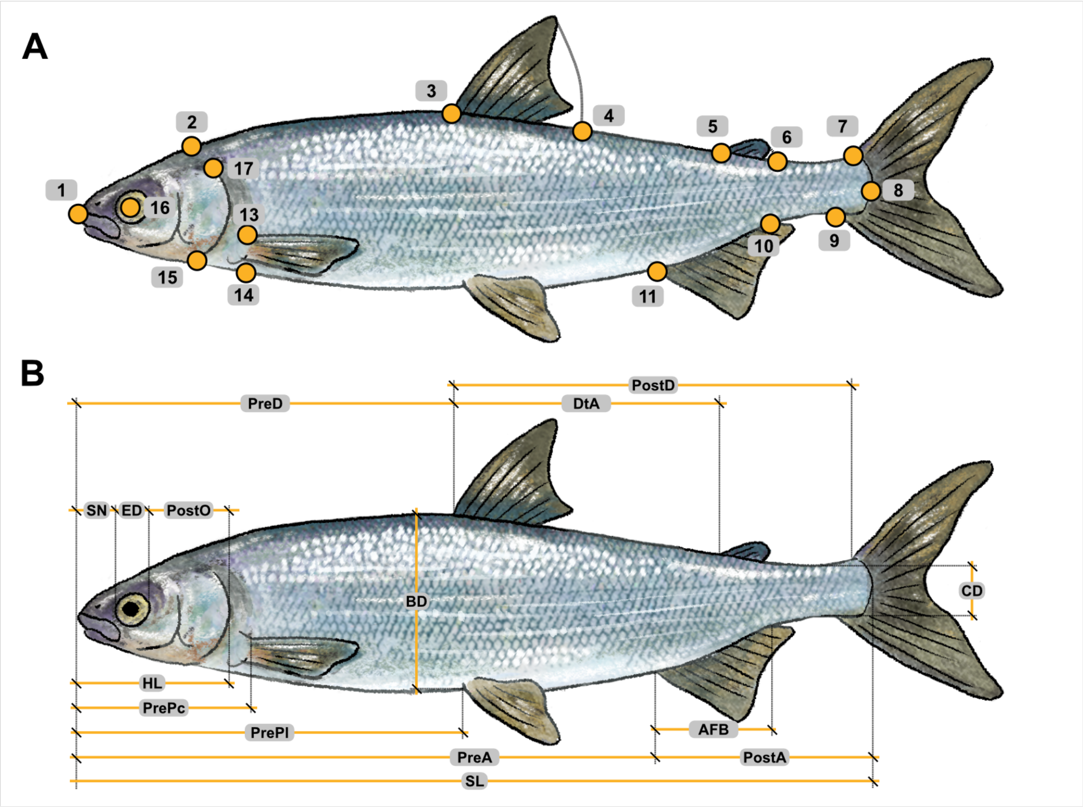


**Figure S1** - Phenotypic analyses. (**A**) Landmarks used for geometric morphometric analyses and (**B**) linear measurements used for morphological analyses. Landmarks and linear measurements are described in Table S5 and S6, respectively.

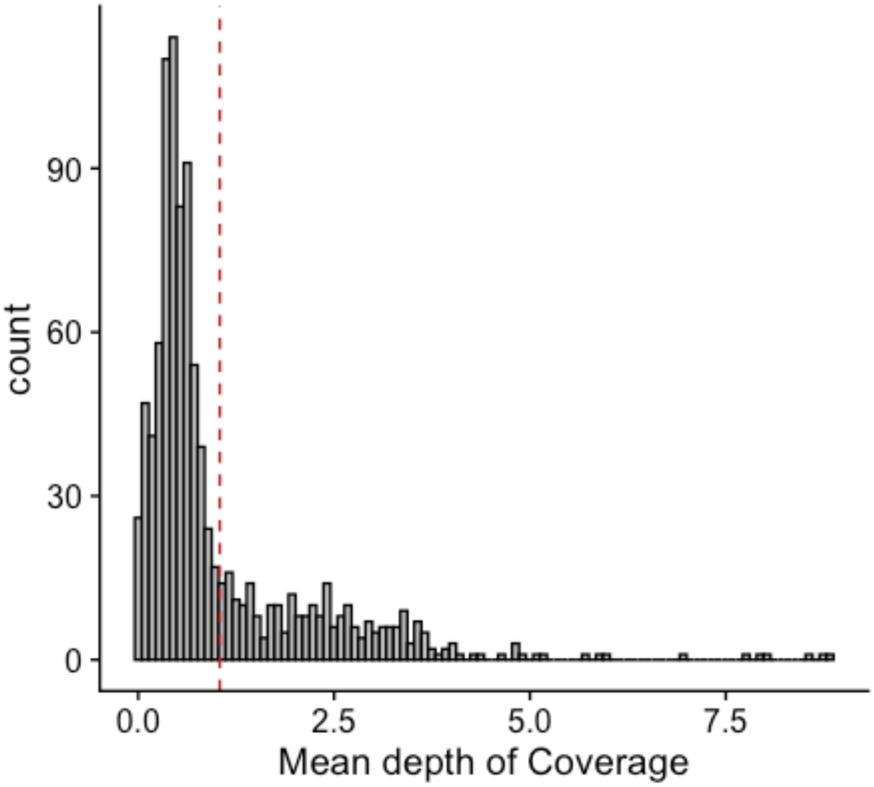


**Figure S2** - Distribution of mean depth of coverage across all sequenced adult and larval individuals. The red line indicates the mean across all individuals (mean = 1.04x).

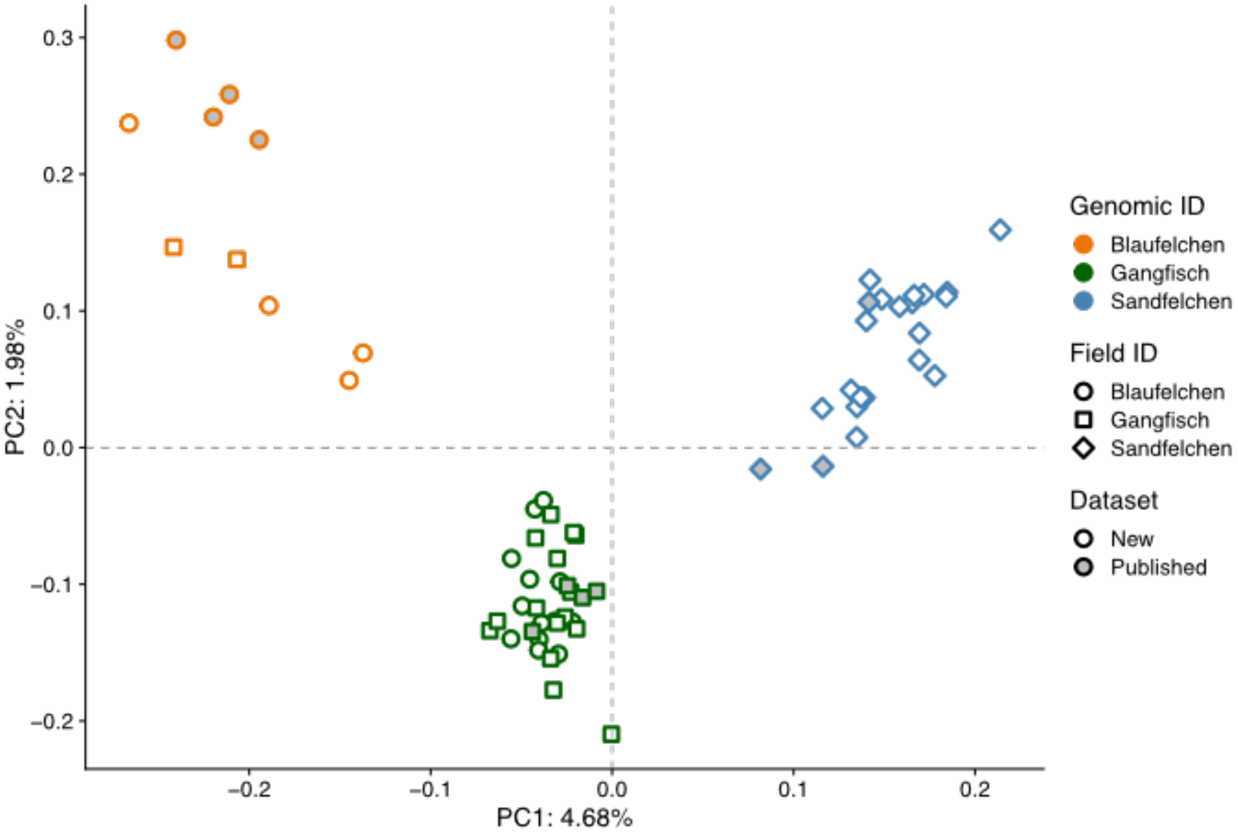


**Figure S3** - Genome-wide PCA for adult whitefish from Upper Lake Constance caught during the targeted fishing campaign (white filled) and published reference samples (grey filled). Individuals are coloured by their genetic species assignment. The identification of species in the field, based on spawning location and morphology, is coded by shape.

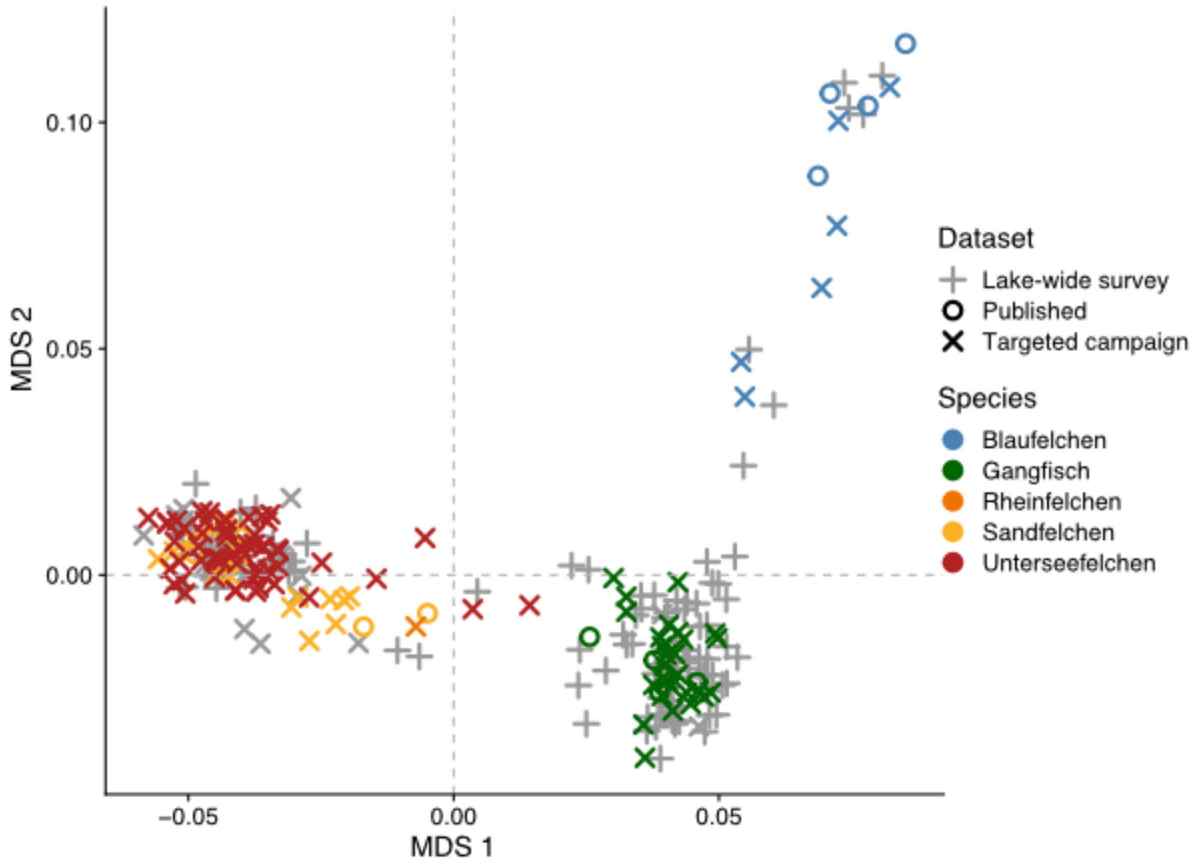


**Figure S4** - Genomic multidimensional scaling plot (MDS) of all adult whitefish, with individual points coloured by their genetic species assignment and datasets highlighted by shape. Lake-wide survey = Individuals caught during the lake-wide fishing campaign in 2024. Targeted campaign = Individuals caught on spawning grounds in 2021-2023. Published = Sequencing data for whitefish derived from the literature (Table S2).

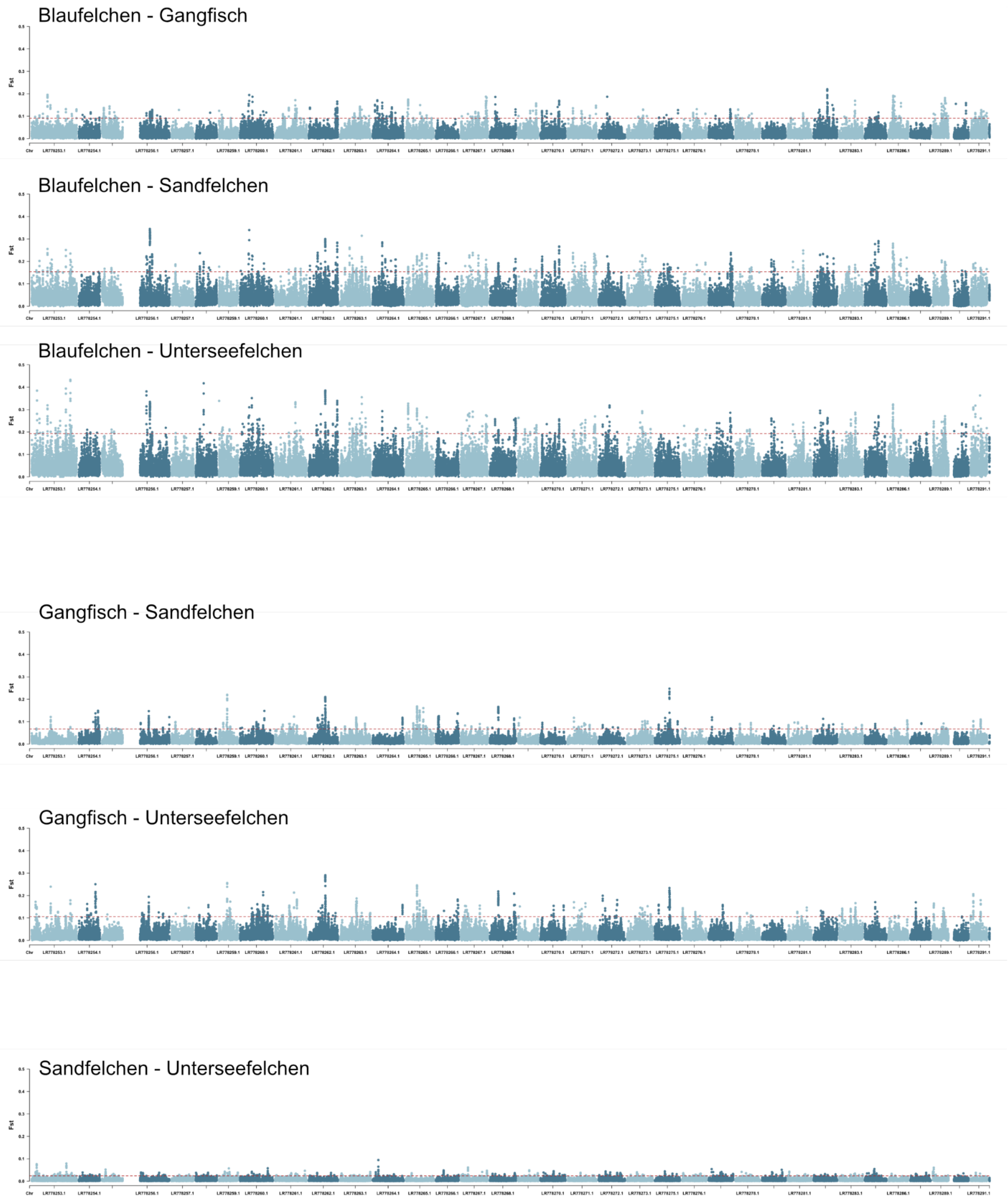


**Figure S5** - Manhattan plots showing Fst genome scans across the entire genome for each species comparison. Fst values are summarised in 50kb sliding windows (25kb steps). Red dotted lines show the 99% percentile of the Fst distribution for each comparison. Chromosomes are shown in alternating colours.

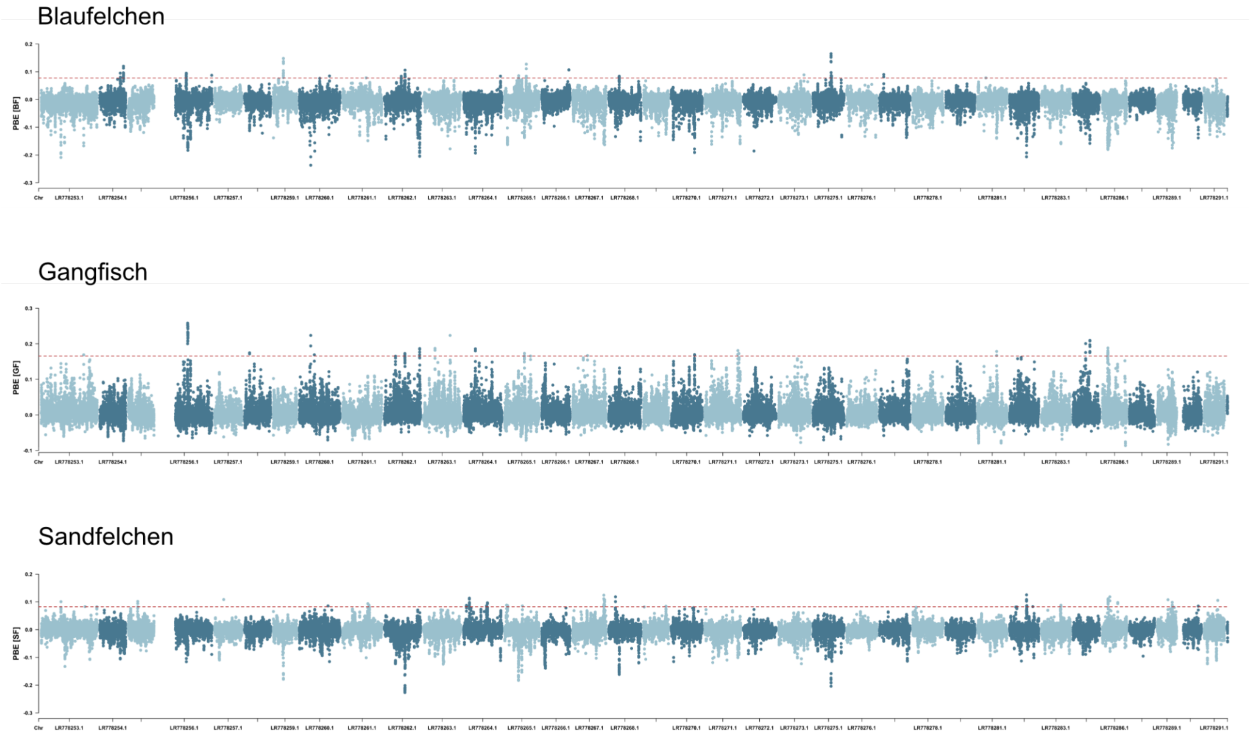


**Figure S6** - Manhattan plots showing population branch excess scores for 50kb sliding windows (25kb steps) across the genome for each species in Upper Lake Constance. Negative PBE values indicate genomic windows for which estimated branch lengths are below the expected value. Red dotted lines indicate the 99.9% percentile of the PBE distribution. Chromosomes are shown in alternating colours.

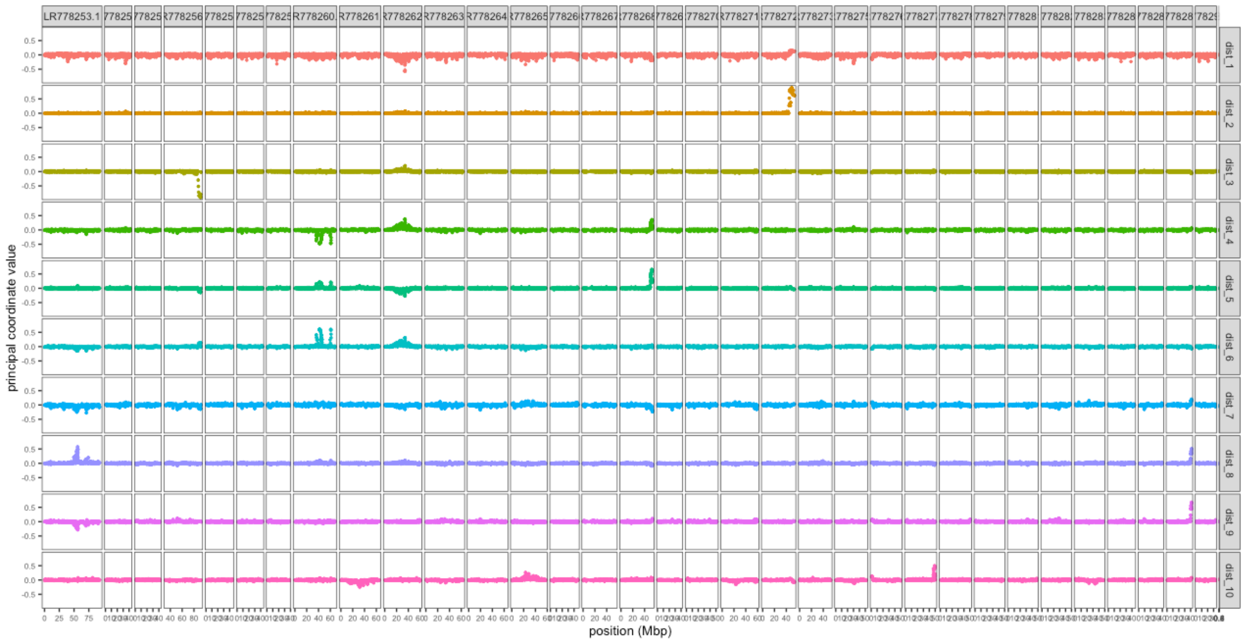


**Figure S7 – LOSTRUCT MDS values by axis.** MDS values across MDS axis 1 to 10 for each 250-SNP window across the genome. Chromosomes are numbered above from 1 to 39.

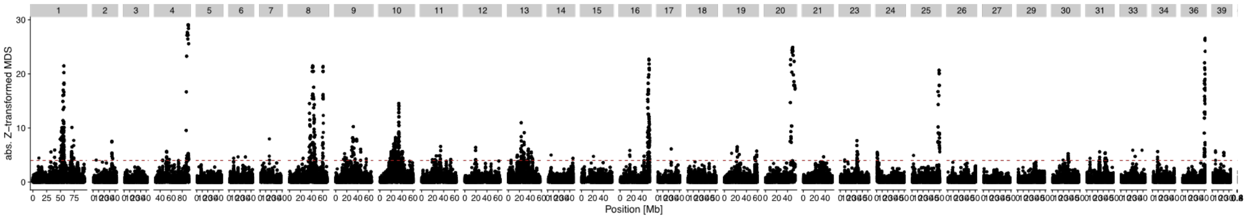


**Figure S8 – LOSTRUCT Manhattan plot.** Manhattan plot showing z-transformed MDS scores for MDS axes 1 to 10 across the genome, with each dot representing a 250-SNP window on one of the MDS axes. The red dotted line shows the threshold of z=4 used for identifying significant outlier windows. Chromosomes are numbered above.

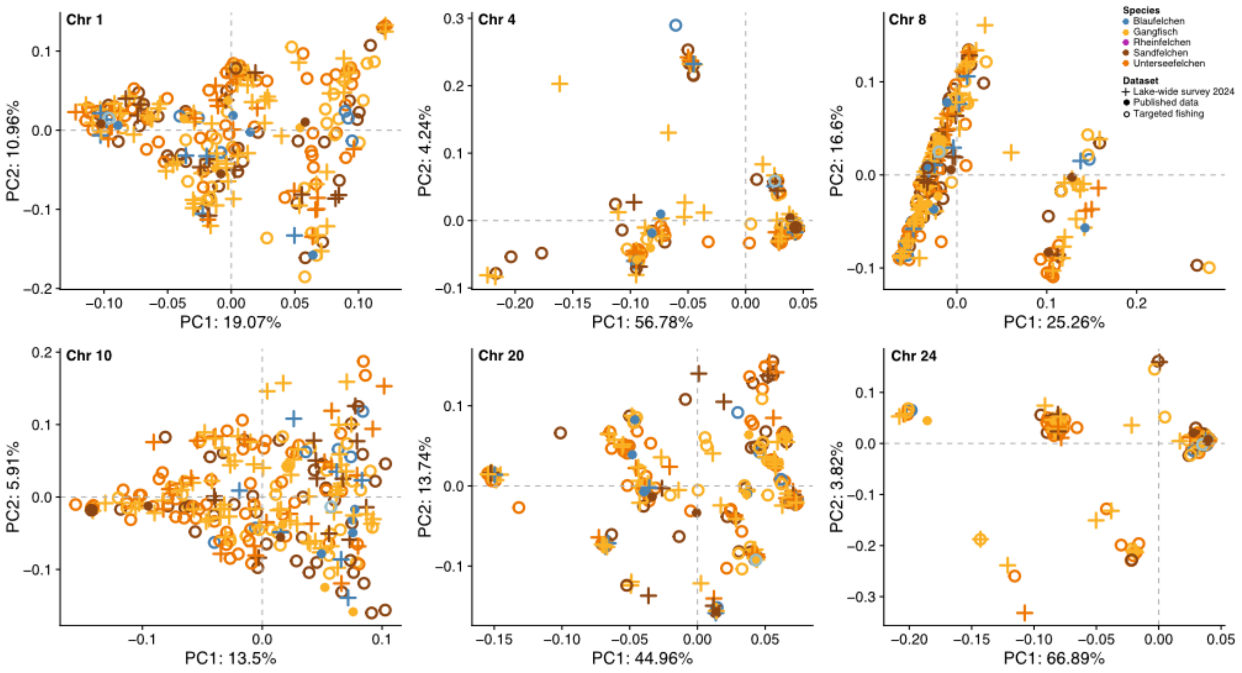


**Figure S9 - LOSTRUCT outliers PCA.** PCA plots (PC1 vs 2) for large genomic regions (>1Mb) with z-transformed MDS values above 4 in the *LOSTRUCT* analysis.

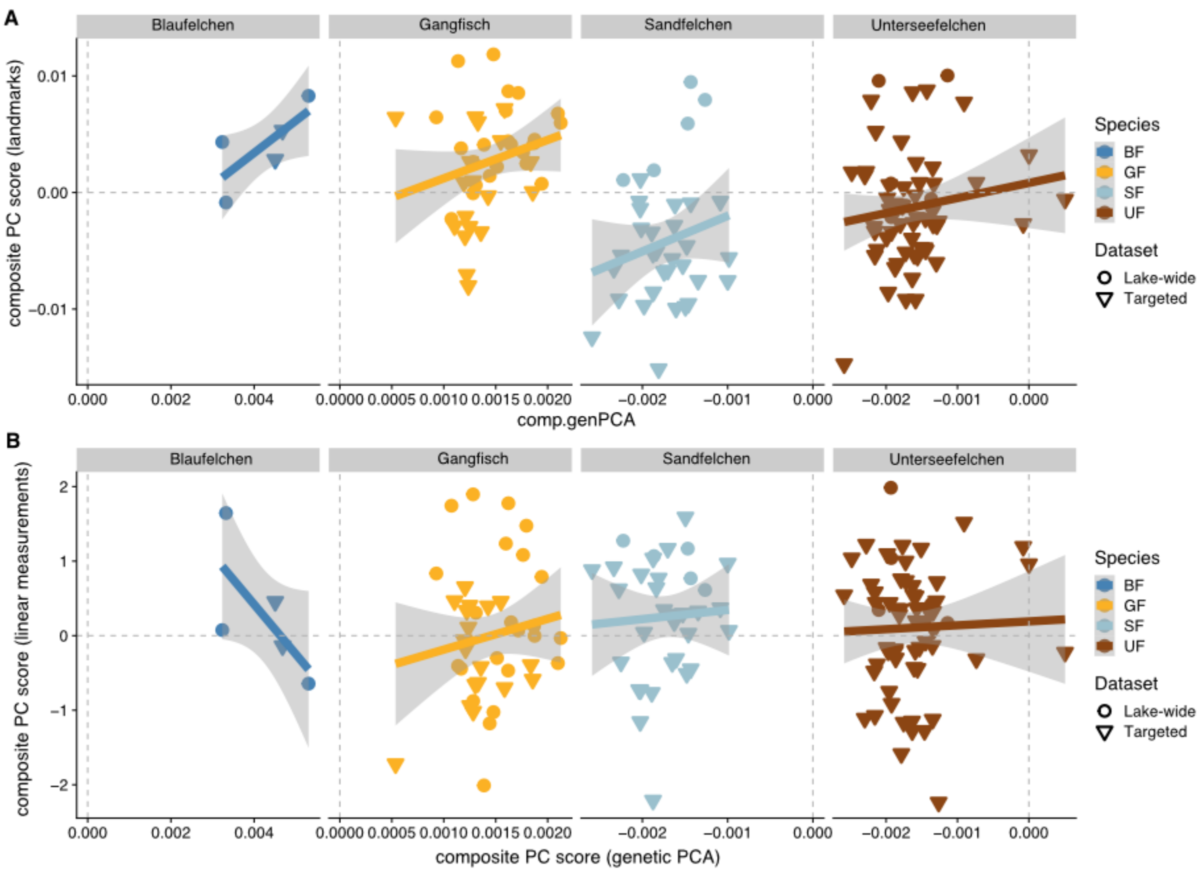


**Figure S10** - Correlations between phenotypic and genetic variation in whitefish by species. Dots are shaped by sampling survey. (A) Shape variation based on landmarks and (B) based on linear measurements.

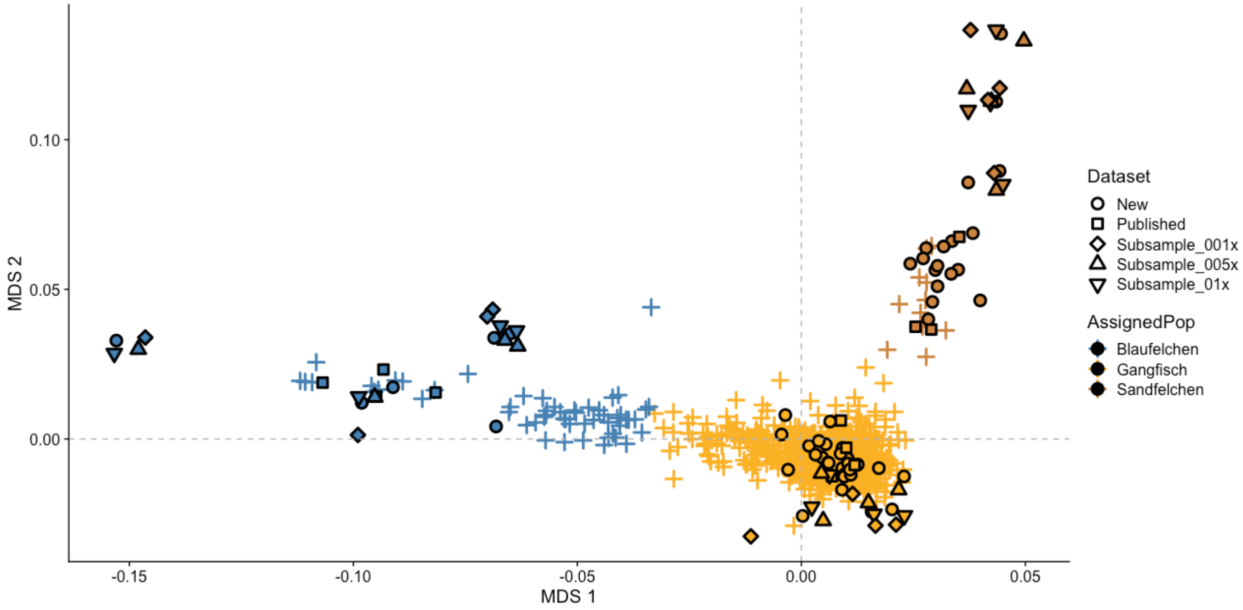


**Figure S11** - MDS plot based on filtered SNPs with larvae (crosses) coloured by species assignment and all adults as circles (newly sequenced data) and squares (published data). Downsampled individuals are shown in different shapes as shown in the legend. Blaufelchen are coloured in blue, Sandfelchen in brown and Gangfisch in yellow.
